# Pseudogene associated recurrent gene fusion in prostate cancer

**DOI:** 10.1101/666933

**Authors:** Balabhadrapatruni VSK Chakravarthi, Pavithra Dedigama-Arachchige, Shannon Carskadon, Shanker Kalyana Sundaram, Jia Li, Kuan-Han Hank Wu, Darshan Shimoga Chandrashekar, James Peabody, Hans Stricker, Clara Hwang, Dhananjay Chitale, Sean Williamson, Nilesh Gupta, Nora M Navone, Craig Rogers, Mani Menon, Sooryanarayana Varambally, Nallasivam Palanisamy

## Abstract

Analysis of next generation transcriptome sequencing data of prostate cancer identified a novel gene fusion formed by the fusion of a protein coding gene (*KLK4*) with a non-coding pseudogene (KLKP1) and expression of its cognate protein. Screening of 659 prostate cancer TMA showed about 32% of positive cases predominantly expressed in higher Gleason grade tumors. Concomitant expression with ERG but not with SPINK1 and other ETS fusion positive tumors. Fusion gene expression potentially regulated by AR and ERG. Antibody specific to the KLK4-KLKP1 fusion protein was validated by immunohistochemistry and western blot methods. Oncogenic properties were validated by *in vitro* and *in vivo* functional studies. Clinical data analysis shows significant association with prostate cancer in young men and overall survival analysis indicate favorable prognosis. Non-invasive detection in urine samples has been confirmed. Taken together, we present a novel biomarker for routine screening of high Gleason grade prostate cancer at diagnosis.

**SIGNIFICANCE:** We discovered and validated a novel prostate cancer (PCa) specific fusion gene involving a protein coding (KLK4) and a pseudogene (KLKP1) and its cognate protein. The unique feature of this fusion gene is the conversion of the noncoding pseudogene into a protein coding gene and its unique expression only in about 30% of high Gleason grade PCa. Expression of this gene is found to be concomitant in ERG fusion positive prostate cancer but mutually exclusive with SPINK1, ETV1, ETV4 and ETV5 positive tumors. Like other ETS family gene fusions, KLK4-KLKP1 can be detected in the urine samples of patients with prostate cancer enabling non-invasive detection of high Gleason grade prostate cancer. Given the unique feature of this fusion oncogenic potential, high Gleason grade specific expression and noninvasive detection, this novel gene fusion has a potential to be used as a biomarker for early detection of high-grade prostate cancer and a therapeutic target.

## INTRODUCTION

Prostate cancer is the most common cancer among men in the United States. Advances in diagnosis, treatment and management has resulted in increased survival rate, yet prostate cancer still remains the second leading cause of cancer-related deaths among American men [1, 2]. One of the major barriers to achieving successful prostate cancer control is the underlying molecular complexity of the disease itself [3]. Morphologically, prostate cancer is well-known to be a diverse disease with patients developing tumors with varying pathological characteristics [4, 5]. Many studies have also indicated that prostate cancer is highly heterogeneous with distinct molecular aberrations observed in patient subgroups [6–8]. For example, roughly 50%-60% of prostate cancer patients are known to carry E26 transformation-specific (ETS) family rearrangements, where *ERG, ETV1, ETV4* or *ETV5* genes fused with androgen regulated 5’ partner genes [9]. Additionally, the overexpression of *SPINK1* has been observed in about 5%-10% of prostate cancer patients [10]. Furthermore 1%-2% of the cases are known to carry *RAF* kinase (*BRAF, RAF1*) gene fusions [11] while the genetic underpinnings in the remaining 30%-40% of the prostate cancer cases are not known [6]. Importantly, distinct molecular changes have been linked with unique disease outcomes [10, 12, 13], indicating complex heterogeneity among patients with respect to disease progression. Therefore, discovery of new molecular markers for further patient stratification and to classify indolent and aggressive prostate cancer is an urgent unmet clinical need to facilitate targeted therapy and effective prostate cancer management.

Currently, prostate cancer diagnosis is primarily based on prostate-specific antigen (PSA) levels and Gleason grade, a scoring system based on the morphology of the prostate tissue [14]. Following the detection of elevated PSA or pro-PSA levels, prostate cancer is identified by the presence of Gleason graded cancer on needle biopsies. The decision to pursue immediate treatment or continue active surveillance is mainly determined using the Gleason grade. However, the rise in PSA is not prostate cancer specific and is multifactorial [15]. Therefore, PSA has been an inadequate diagnostic marker, in some cases leading to overdiagnosis and unnecessary treatment. Though high Gleason grade tumors are known to be clinically aggressive, whether low Gleason grade tumors require treatment has been debated [16]. While intervention in low Gleason grade cancers may result in overtreatment, watchful waiting may also pose an unnecessary risk and additional burden of repeat biopsies. Given these limitations of the existing markers and the recognition of prostate cancer as a heterogeneous disease, molecular markers specific to distinct patient subgroups, are required as alternatives for both initial cancer diagnosis and distinguishing aggressive cancer from indolent disease.

Although several recurrent molecular alterations have been identified in a subset of prostate cancer cases, the genetic aberrations in prostate cancer patients negative for all the known molecular makers remain to be studied. Moreover, most prostate cancer molecular studies have been carried out on Caucasian American patients with little representation from the African American (AA) population [17]. Given the unique ancestral background of AAs and the aggressive nature of prostate cancer, the genetic underpinnings to understand the racial disparity in the incidence of prostate cancer markers is not well studied. Therefore, the study of additional molecular aberrations using large cohorts of a racially diverse population is a pressing need in prostate cancer research. In addition to identifying subtype specific prostate cancer diagnostic and prognostic markers, such studies may also facilitate the development of novel therapeutic approaches by uncovering molecular alterations, which may be pharmacologically targeted in distinct patient subgroups.

Given the need for identifying novel molecular markers in prostate cancer patients, we investigated the expression patterns of pseudogenes in 89 prostate cancer patient samples using a paired-end next generation sequencing approach [18]. Often considered as dysfunctional relatives of known protein-coding genes, pseudogenes have recently been implicated in cancer with roles in gene regulation [19]. While we observed distinct expression changes in several pseudogenes in prostate cancer compared to normal prostate tissue, we also noted the rare occurrence of a chimeric transcript formed through the fusion of the androgen regulated gene *KLK4* (Kallikrein Related Peptidase 4) with the adjacent pseudogene *KLKP*1 (Kallikrein Pseudogene 1). Importantly, the fusion converts the *KLKP1* pseudogene to a protein-coding gene with a predicted chimeric protein of 164 amino acids, of which 55 amino acids are derived from the pseudogene part due to a shift in the open reading frame of the fusion formed by trans-splicing mechanism rather than chromosomal rearrangement [18]. Although a few pseudogenes have been previously reported to be expressed as proteins [20, 21], *KLK4-KLKP1* is a rare example where gene fusion leading to conversion of a non-coding pseudogene to a protein-coding gene. Further studies showed that *KLK4-KLKP1* fusion is both prostate tissue and cancer specific, suggesting a possible role in prostate cancer formation [18]. Both the prostate cancer specific expression and the intriguing nature of the *KLK4-KLKP1* fusion warrant further functional studies to understand the role of *KLK4-KLKP1* in prostate cancer development. Therefore, in this study, we explored the prevalence, the expression pattern, noninvasive detection, and the oncogenic properties of *KLK4-KLKP1* to investigate the potential of *KLK4-KLKP1* fusion gene as a novel molecular marker in prostate cancer.

## RESULTS

Both *KLK4* and *KLKP1* belong to the kallikrein family of serine proteases, a cluster of genes located on chromosome 19(q13.33-q13.41). The gene cluster contains 15 members including *KLK3*, which is commonly known as PSA [22]. The *KLK4-KLKP1* fusion is formed by a trans-splicing mechanism or an in-frame fusion due to a microdeletion of the region between the adjacent genes, *KLK4* and *KLKP1*, leading to the fusion of the first 2 exons of *KLK4* with exon 4 and 5 of *KLKP1* (Fig. 1A **and Supplementary Fig. S1,** GenBank ID 2227664). The resulting chimeric sequence predicts a 164 amino acid protein, of which 55 amino acids are derived from KLKP1 (Fig. 1B). According to data on the GTEx portal, full length KLKP1 is exclusively expressed in normal prostate tissue (**Supplementary Fig. S2**). In contrast, quantitative PCR (qRT-PCR) analysis of prostate cancer samples, prostate cell lines, benign prostate tissues and other solid cancers revealed that *KLK4-KLKP1* fusion transcript is prostate cancer specific and expressed in a subset of cases [18]. However, the study included only a limited number of prostate cancer samples (n = 36) and the occurrence of *KLK4-KLKP1* in a large, racially inclusive cohort must be explored to determine the prevalence of *KLK4-KLKP1* in the prostate cancer patient population. Therefore, we studied the expression of *KLK4-KLKP1* on a larger patient cohort using fusion transcript specific anti-sense oligonucleotide probe by RNA *in situ* hybridization (RNA-ISH). Specifically, we constructed tissue microarrays (TMAs) using prostate cancer tissues obtained from 659 radical prostatectomy (RP) specimens at the Henry Ford Health Systems. The cohort was racially inclusive with 380 Caucasian, 250 AA and 29 patients belonging to other racial groups. Each TMA contained 3 cores obtained from different regions of the RP prostate from each patient (**Supplementary Fig. S3**). The individual tissue cores in each patient were reviewed and the highest tumor grade observed was assigned to each case. Thus the TMAs included 612 patient cases with all cores carrying prostate cancer (Gleason grade group 1 [3 + 3 = 6] - 110, Gleason grade group 2 [3 + 4 = 7] - 247, Gleason grade group 3 [4 + 3 = 7] - 119, Gleason grade group 4 [4 + 4 = 8] - 94, and Gleason grade group 5 [4 + 5 = 9; 5 + 4 = 9 and 5 + 5 = 10] - 42). The rest of the cases consisted of 23 cases with benign, 21 cases with high grade prostate intraepithelial neoplasia, 2 cases with stroma and 1 case with atypical cores. RNA-ISH was carried out using an antisense RNA probe specific to the *KLK4-KLKP1* fusion. The TMA slides were then reviewed for the intensity of the RNA-ISH signal. A score of expression ranging from 0 to 4+ was given according to the intensity of the RNA-ISH signal where, 0 indicated no detectable RNA-ISH signal, while 4+ was assigned to the highest level of RNA-ISH signal [23].

**Figure 1:**
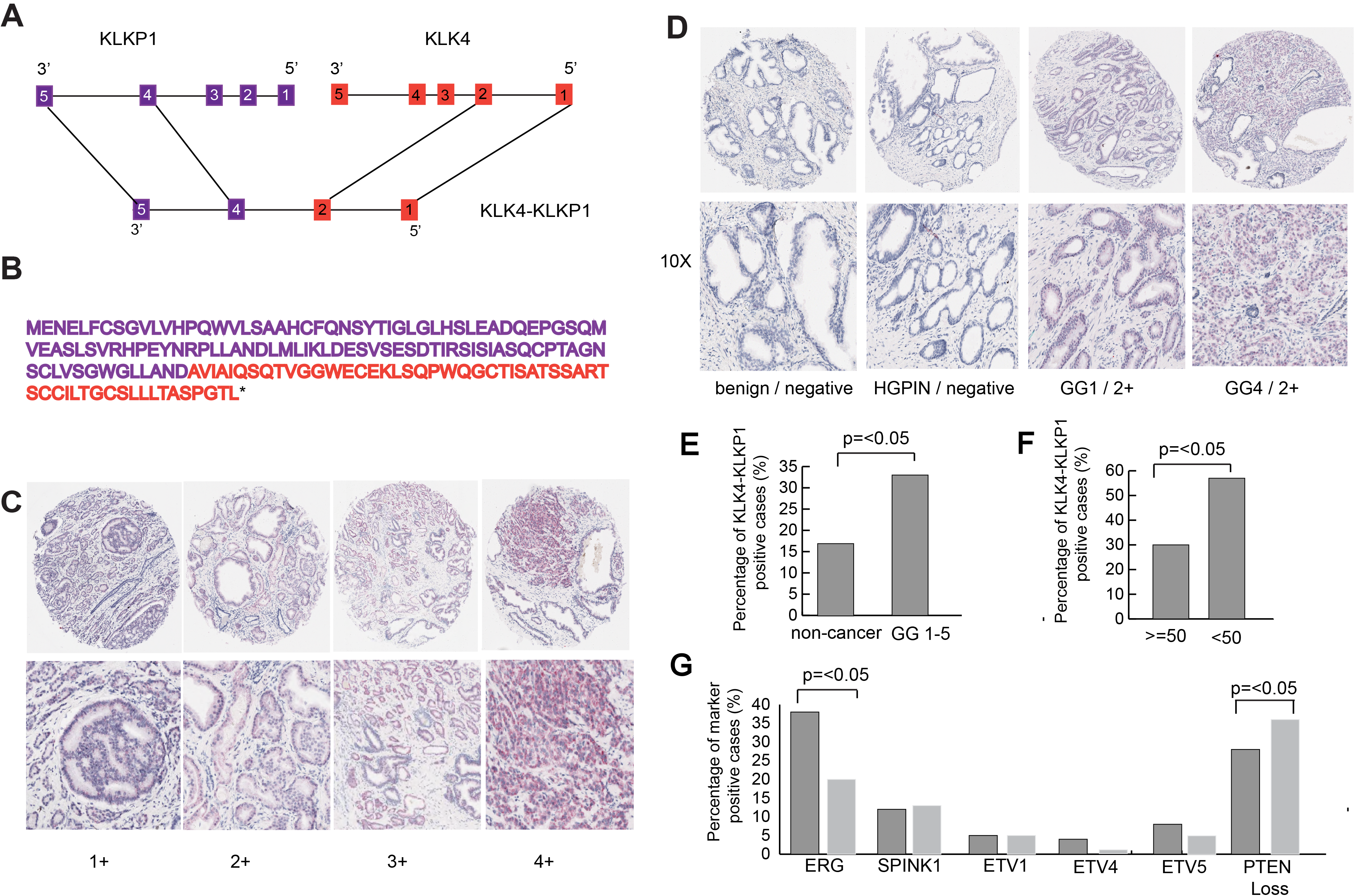
The structure of *KLK4-KLKP1* fusion and the RNA-ISH screening of *KLK4-KLKP1* in tissue micro arrays. (A) Schematic diagram of the structure of *KLK4-KLKP1* fusion. *KLK4-KLKP1* is formed through the fusion of exon 1 and 2 of *KLK4* gene with exon 4 and 5 of *KLKP1*. (B) The predicted sequence of *KLK4-KLKP1* fusion protein. The sequence in purple is derived from KLK4 while the sequence in red is originating from KLKP1. (C) The expression of *KLK4-KLKP1* in prostate tissue cores detected by RNA-ISH. The bottom set of images show an enlarged section of the corresponding tissue core in the top set of images. 1+ to 4+ indicate the intensity of *KLK4-KLKP1* RNA-ISH staining. (D) Prostate cancer specific expression of *KLK4-KLKP1*. *KLK4-KLKP1* RNA-ISH staining in benign, HGPIN and prostate cancer tumor cores are shown. The bottom set of images contains a magnified area of the images on the top. 1+ to 4+ refer to the intensity of the *KLK4-KLKP1* RNA-ISH staining. (E) *KLK4-KLKP1* is expressed more in the prostate cancer patients (GG1-5) compared to non-cancer (benign, HGPIN, atypical and stroma) cases. The percentage of cases showing a positive *KLK4-KLKP1* RNA-ISH signal among non-cancer and GG1-5 groups is shown. P-value was calculated based on Pearson’s chi-square test. (F) *KLK4-KLKP1* is expressed more in young patients. The percentages of cases with positive *KLK4-KLKP1* RNA-ISH signal in the young patient (age lower than 50 years) and old patient groups (age equal to or higher than 50 years) are shown. P-value was calculated based on Pearson’s chi-square test. (G) *KLK4-KLKP1* expression is associated with *ERG* overexpression. *SPINK1*, *ETV1*, *ETV4* and *ETV5* overexpression is mutual from *KLK4-KLKP1* expression. *PTEN* loss is significantly lower in cases with *KLK4-KLKP1* expression. The percentages of cases showing positive signal for *ERG*, *SPINK1*, *ETV1*, *ETV4*, *ETV5* or *PTEN* loss among *KLK4-KLKP1* RNA-ISH positive cases (dark grey bars) and *KLK4-KLKP1* RNA-ISH negative cases (light grey bars) are shown. P-value was calculated based on Pearson’s chi-square test. Abbreviations: GG, Gleason grade; HGPIN, high grade prostate intraepithelial neoplasia; ISH, in situ hybridization.

Of the 659 cases in the cohort, 209 (32%) were positive for *KLK4-KLKP1* fusion, indicating the recurrent nature of *KLK4-KLKP1* among prostate cancer patients. Most of the *KLK4-KLKP1* positive cases showed RNA-ISH signal intensity of 1+ (130 cases; Fig. 1C) while more intense RNA-ISH signal 2+ was observed in 66 cases, 3+ in 12 cases and 4+in 1 case (Fig. 1C) and remaining cases were “0” or negative, suggesting varying expression levels among patients. To further confirm that KLK4-KLKP1 is specific to prostate cancer, we then explored the association of *KLK4-KLKP1* RNA-ISH signal with Gleason grade by using Pearson’s chi-square test. The results showed that *KLK4-KLKP1* is exclusively expressed in prostate cancer tissues compared to benign, high grade prostate intraepithelial neoplasia and atypical prostate tissues (Fig. 1D, Fig. 1E, **and Table 1**), confirming that *KLK4-KLKP1* expression is prostate cancer specific. Additionally, we also analyzed if KLK4-KLKP1 expression is associated with Gleason grade group and found no associations with distinct Gleason grade groups (**Table 2**).

Next, we investigated if *KLK4-KLKP1* fusion displays racial disparity in the incidence. The 209 positive cases included 128 Caucasian Americans (34%), 69 AAs (28%) and 12 patients from other races (41.4%). Pearson’s chi-square test analysis revealed that prevalence of KLK4-KLKP1 is high in Caucasian American compared with AA patients but not statistically significant (**Table 3 and Supplementary Fig. S4**), demonstrating no racial bias in the incidence. We also explored if *KLK4-KLKP1* expression is related with patient age. We categorized the patients into 2 groups as young (age ranging from 40 to 50 years) and old (age ranging from 51 to 83 years). Pearson’s chi-square test showed significantly higher expression of *KLK4-KLKP1* in young age group compared to the old age group (Fig. 1F **and Table 4**).

The other common prostate cancer specific mutations such as ETS gene fusions and *SPINK1* overexpression are known to occur in a mutually exclusive manner. Therefore, we also analyzed the association of *KLK4-KLKP1* fusion expression with ETS gene fusions and *SPINK1* expression. We screened the same set of TMAs by using dual immunohistochemistry (IHC) for *ERG* and *SPINK1* and dual RNA-ISH for *ETV1, ETV4* and *ETV5*. By using Pearson’s chi-square test, we observed that *KLK4-KLKP1* expression is associated with *ERG*+ cases (Fig. 1G, **Supplementary Fig. S5, and Table 1**). However, no such association was observed with *SPINK1*, *ETV1*, *ETV4* and *ETV5* (Fig. 1G **and Table 5)**, suggesting concurrent expression of *KLK4-KLKP1* with distinct ETS gene fusion positive cases. Next, we investigated if *KLK4-KLKP1* is related with *PTEN* loss, another common prostate cancer mutation that is associated ERG+ and aggressive disease [24–26]. We carried out IHC for *PTEN* on the same set of TMAs and found that *PTEN* deletion was significantly lower in *KLK4-KLKP1* positive cases compared to *KLK4-KLKP1* negative cases (Fig. 1G **and Table 1**). Given that *ERG* is known to co-occur with *PTEN* loss [27], we further analyzed if there is any significant difference in *PTEN* loss in cases showing both *ERG* fusion and *KLK4-KLKP1* compared to the rest of the cases and found no significant difference in *PTEN* status in cases with *ERG* fusion and *KLK4-KLKP1* expression, suggesting that *KLK4-KLKP1* may represent a distinct subtype of prostate cancer.

Having thus confirmed the recurrent and the prostate cancer specific occurrence of *KLK4-KLKP1* fusion, we then studied the expression of *KLK4-KLKP1* fusion protein. Based on the sequence, the *KLK4-KLKP1* fusion gene is predicted to generate a full-length protein of 164 amino acids of which 55 are derived from the KLKP1 pseudogene (Fig. 1B). To validate the *KLK4-KLKP1* expression as a full-length protein, we generated adenoviral constructs carrying the N-FLAG-tagged *KLK4-KLKP1* fusion gene and transfected HEK293 cells. To stabilize the protein levels of *KLK4-KLKP1*, the cells were treated with the proteasome inhibitor bortezomib. As a control, bortezomib treated cells transfected with vector DNA alone were used. Expression of the fusion transcript was confirmed by qRT-PCR using fusion specific primers (Fig. 2A). Cell lysates were analyzed by western blotting using an anti-N-FLAG antibody. Importantly, we observed a FLAG-specific protein band around 17kDA (Fig. 2B), confirming the expression of *KLK4-KLKP1* as a full-length protein. For additional validation, we also checked the expression of N-FLAG-tagged *KLK4-KLKP1* using the anti-FLAG antibody in the normal prostate cell line RWPE-1, transfected with and without N-FLAG-tagged *KLK4-KLKP1* adenovirus construct. Notably, we detected anti-FLAG specific protein band only in the transfected RWPE1 cells (**Supplementary Fig. S6**). Furthermore, we also developed a *KLK4-KLKP1* specific polyclonal antibody (Eurogentech, Seraing, Belgium) using the antigenic peptide “CTISATSSARTS” (Fig. 1B) derived from the *KLKP1* pseudogene region of the fusion protein. After cell lysis and SDS-PAGE, we probed HEK293 lysates transfected with and without N-FLAG-tagged *KLK4-KLKP1* adenovirus construct with the *KLK4-KLKP1* specific antibody using western blot. A protein band around 17kDA was observed further confirming the expression of the chimeric *KLK4-KLKP1* protein (Fig. 2B, 2E) and the specificity of the antibody to the fusion protein.

**Figure 2:**
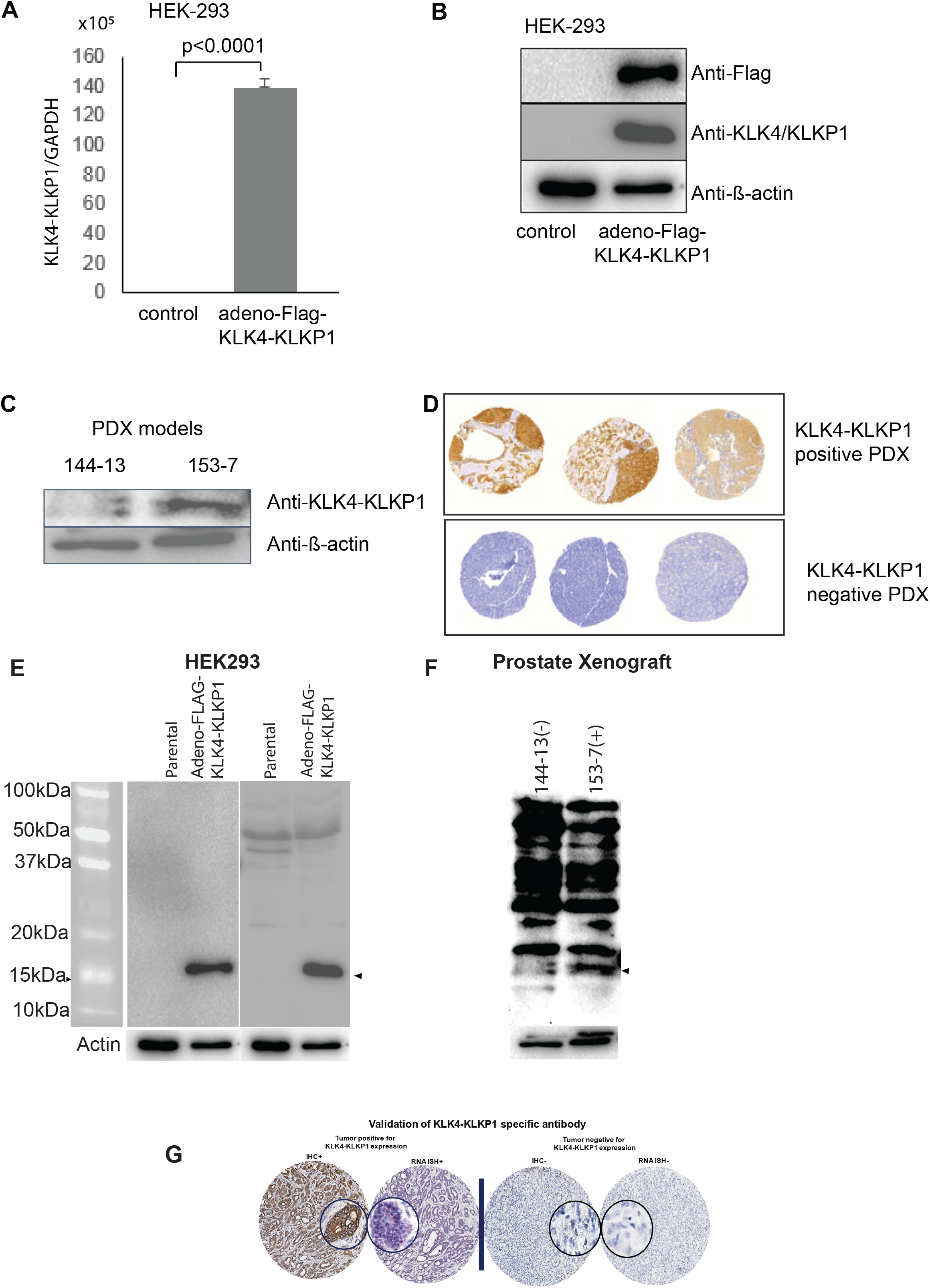
Validation of the expression of *KLK4-KLKP1* protein in HEK-293 cells and PDX tissues. (A) The qRT-PCR analysis HEK-293 cells transfected with and without FLAG tagged-*KLK4-KLKP1*. HEK-293 cells were transfected with adenoviral vectors carrying FLAG tagged-*KLK4-KLKP1* (adeno-FLAG-*KLK4-KLKP1*). As a control untransfected cells treated with bortezomib were used. The expression of *KLK4-KLKP1* was confirmed by qRT-PCR. (B) Western blot analysis of HEK-293 cells transfected with FLAG-tagged *KLK4-KLKP1* using anti-FLAG, anti-*KLK4-KLKP1* and anti-β-actin antibody. (C) Western blot analysis of *KLK4-KLKP1* qRT-PCR negative (MDA PCa144-13) and qRT-PCR positive (MDA PCa 153-7) PDX tissues using anti-*KLK4-KLKP1* and anti-β-actin antibody. (D) IHC staining of KLK4-KLKP1 qRT-PCR positive and qRT--PCR negative PDX models. Abbreviations: PDX, patient derived xenografts; qRT-PCR, quantitative PCR. Images of original western blots show anti-N-FLAG antibody (2E-left), and anti-KLK4-KLKP1antibody (2E-right). Images of original western blots show, lysates from the prostate xenografts each positive (MDA PCa 153-7) and negative (MDA PCa 144-13) for endogenous expression of KLK4-KLKP1 transcript were probes with anti-KLK4-KLKP1 antibody (2F). Validation of KLK4-KLKP1 specific antibody in comparison with RNA-ISH (2G). PCa tissue confirmed to be positive by RNA-ISH (left) is positive for the antibody whereas the tumor negative for KLK4-KLKP1 by RNA-ISH also negative for the antibody by IHC, thus confirming the specificity of the new antibody to the KLK4-KLKP1 fusion protein.

In order to assess the expression of *KLK4-KLKP1* in metastatic prostate cancer, we then analyzed the expression of *KLK4-KLKP1* in prostate cancer patient derived xenografts (PDX)[28]. We first screened the expression of *KLK4-KLKP1* using qRT-PCR and identified 17 out of 31 PDX models positive for endogenous expression of *KLK4-KLKP1* (**Supplementary Fig. S7; Table 6**). Then, we selected one of the PDX tissues (MDA PCa 153-7) expressing high levels of *KLK4-KLKP1* and one with no detectable levels of *KLK4-KLKP1* (MDA PCA 144-13). After protein isolation and separation on SDS-PAGE, the lysates were probed with the *KLK4-KLKP1* specific antibody using western blot. Importantly, we observed a protein band around 17kDA only in the *KLK4-KLKP1* positive PDX (Fig. 2C, 2F), indicating the endogenous expression of *KLK4-KLKP1* fusion protein in metastatic prostate cancer patients. Additionally, we also screened the expression of *KLK4-KLKP1* in xenograft tissues using IHC with the *KLK4-KLKP1* specific antibody. While *KLK4-KLKP1* expression was observed in qRT-PCR positive PDX tissues, minimal or no *KLK4-KLKP1* IHC signal was seen in qRT-PCR negative xenografts (Fig. 2D), further suggesting the presence of *KLK4-KLKP1* protein in a subset of prostate cancer patients. Comparison of IHC results with RNA-ISH positive and negative tissue showed specificity of the antibody to KLK4-KLKP1 RNA-ISH positive tissue only (Fig.2G).

Given the exclusive expression of *KLK4-KLKP1* in prostate cancer, next we explored the functions of *KLK4-KLKP1* by studying the oncogenic properties of the fusion gene. Specifically, we established RWPE-1 cells with stable expression of *KLK4-KLKP1* by transfection with lentiviral constructs carrying FLAG-tagged *KLK4-KLKP1*. As controls, cells stably transfected with a LACZ control (LACZ) and un-transfected RWPE-1 cells were used. We first confirmed the expression of *KLK4-KLKP1* by qRT-PCR. The results showed significant expression of *KLK4-KLKP1* in transfected cells compared to both the un-transfected cells and the LACZ control (Fig. 3A). Then we investigated the effect of *KLK4-KLKP1* on cell proliferation by measuring the number of cells using a Coulter particle counter. Compared to the un-transfected cells and the LACZ control, a notable increase in the cell number was seen over time in *KLK4-KLKP1* transfected cells (Fig. 3B), indicating a role of *KLK4-KLKP1* on cell proliferation. Next, we studied the effect of *KLK4-KLKP1* in cell invasion using the Matrigel invasion assay. Importantly, a significant increase in the number of invaded cells was observed with *KLK4-KLKP1* transfected cells compared to both the un-transfected and the LACZ control (Fig. 3C). For additional validation, we also transiently transfected PrEC, another normal prostate cell line with *KLK4-KLKP1*. As controls, un-transfected cells and cells transfected with a LACZ control were used. Additionally, we also used cells transfected with *EZH2*, which has been shown to increase invasion of prostate cancer and other cancer cells [29, 30] as a positive control. The invasion of cells was then examined by the Matrigel invasion assay. Like RWPE-1, PrEC cells also showed a significant increase in the number of invaded cells compared to both the un-transfected and the LACZ control (Fig. 3D). As expected, cells transfected with EZH2 also demonstrated increased invasion compared to the LACZ control and the un-transfected cells (Fig. 3D). In all, our studies indicate that *KLK4-KLKP1* promote both cell proliferation and invasion of prostate cells, suggesting an oncogenic role for *KLK4-KLKP1* fusion.

**Figure 3:**
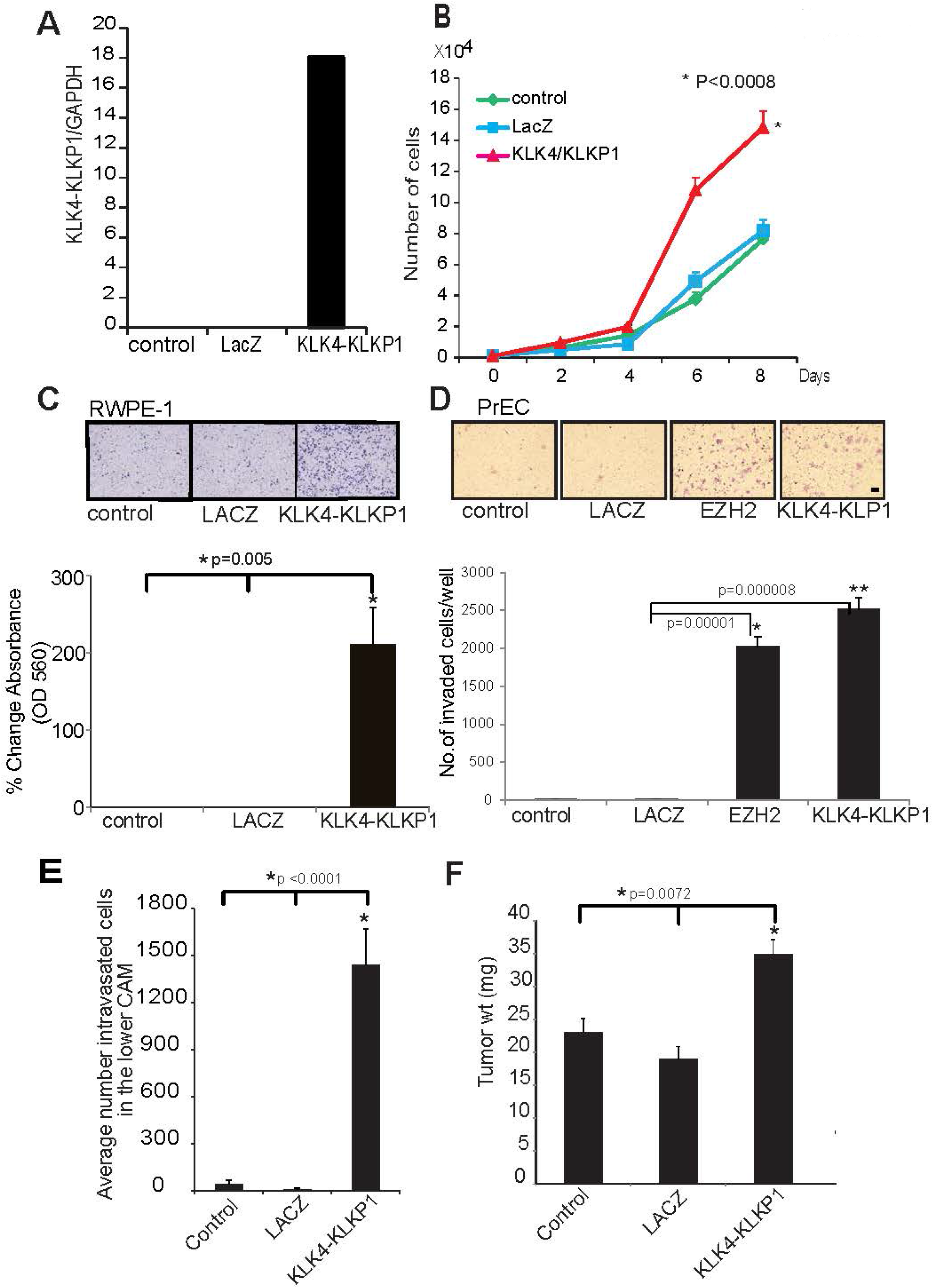
Functional characterization of *KLK4-KLKP1*. (A) qRT-PCR validation of *KLK4-KLKP1* expression in RWPE-1 cells after stable transfection with FLAG tagged-KLK4-KLKP1. As controls untransfected cells (control) and cells transfected with LacZ were used. (B) Analysis of cellular proliferation in RWPE-1 cells stably expressing FLAG tagged *KLK4-KLKP1*. Cells were plated in 96-well plates. The number of cells was measured on days 2, 4, 6 and 8 using a Coulter particle counter. Cells untransfected and transfected with LACZ were used as controls. (C) Analysis of cell invasion in RWPE-1 cells. The invasion of RWPE-1 cells stably transfected with either FLAG tagged-*KLK4-KLKP1* or LacZ was studied using the Boyden chamber assay. Untransfected cells were also used as a control. After invasion of cells into the invasion chamber, cells were fixed and visualized using crystal violet. Additionally, the invasion chamber membranes carrying the fixed cells were dipped in glacial acetic acid and the absorbance at 560 nm was also measured. Representative images of the crystal violet stained cells that underwent invasion in each case and the absorbance at 560 nm are shown. (D) Analysis of cell invasion in PrEC cells. The cellular invasion in PrEC cells transfected with FLAG tagged-*KLK4-KLKP1* was performed as described in Figure 2C. The number of invaded cells were counted and plotted. In addition to LACZ and untransfected cells, PrEC cells transfected with *EZH2* were also used a control. (E) Intravasation of RWPE-1 cells measured using CAM assay. RWPE-1 cells stably transfected with FLAG tagged-*KLK4-KLKP1*, were implanted on eggs. The presence of intravasated cells in the lower CAM was assessed by quantitative human Alu-specific PCR. Untransfected cells and cells transfected with LACZ were used as controls. (F) Analysis of weight of extraembryonic tumors isolated from eggs implanted with RWPE-1 cells stably expressing FLAG-tagged *KLK4-KLKP1*. Cells transfected with LACZ and untransfected cells were used as controls. Abbreviations: CAM, chicken chorioallantoic membrane.

In order to further understand the oncogenic properties of *KLK4-KLKP1*, we also studied the effects of *KLK4-KLKP1* fusion on intravasation and tumor formation using the chicken chorioallantoic membrane (CAM) *in vivo* assay [31, 32]. We implanted eggs with RWPE-1 cells stably expressing *KLK4-KLKP1* and then checked for the presence of intravasated cells in the lower CAM by using quantitative human Alu-specific PCR. As controls, eggs implanted with either un-transfected cells or cells stably transfected with a LACZ control were used. Notably, we observed a marked intravasation by *KLK4-KLKP1* transfected cells in the lower CAM compared to both un-transfected cells and LACZ control (Fig. 3E). Additionally, we also isolated and weighed the extraembryonic tumors from eggs implanted with either *KLK4-KLKP1* transfected cells or controls. The tumors isolated from eggs implanted with cells expressing *KLK4-KLKP1* showed significantly higher weight than the tumors isolated from eggs treated with the un-transfected cells and the LACZ control (Fig. 3F). Overall, the results establish that *KLK4-KLKP1* drives intravasation and tumor formation in prostate cells, indicating a potential role in prostate cancer development.

Further, we investigated the molecular mechanisms underlying the oncogenic functions of *KLK4-KLKP1* fusion. We conducted a gene expression microarray analysis using RWPE-1 cells stably transfected with *KLK4-KLKP1*. As the control, cells transfected with LACZ control were used. After RNA isolation, and microarray analysis, we observed a significant number of genes expressed differently between the RWPE-1 cells transfected with *KLK4-KLKP1* and the LACZ control. We selected the genes showing a fold change value of more than 1 in 2 independent replicates and generated a heat map with the top 100 genes differentially expressed (Fig. 4A). We noted genes both upregulated and downregulated in cells expressing the *KLK4-KLKP1* fusion, suggesting a possible function for *KLK4-KLKP1* in gene expression regulation. Further, we also carried out a gene set enrichment analysis [33] to explore any overlap between the differentially expressed genes observed with *KLK4-KLKP1* transfection and other curated gene sets. Importantly, we noted enrichment of 2 curated gene sets, one involving genes upregulated in endometroid endometrial metastatic tumor and the other containing genes overexpressed in melanoma metastatic cancer (Fig. 4B), indicating that the genes affected by *KLK4-KLKP1* are associated with metastatic cancer. As a further step, we also carried out a KEGG pathway analysis using the DAVID tool [34]. The genes differentially affected by *KLK4-KLKP1* were shown to be associated with several cancer-related pathways (Fig. 4C), further implying that *KLK4-KLKP1* may regulate the expression of genes involved in cancer and metastasis.

**Figure 4:**
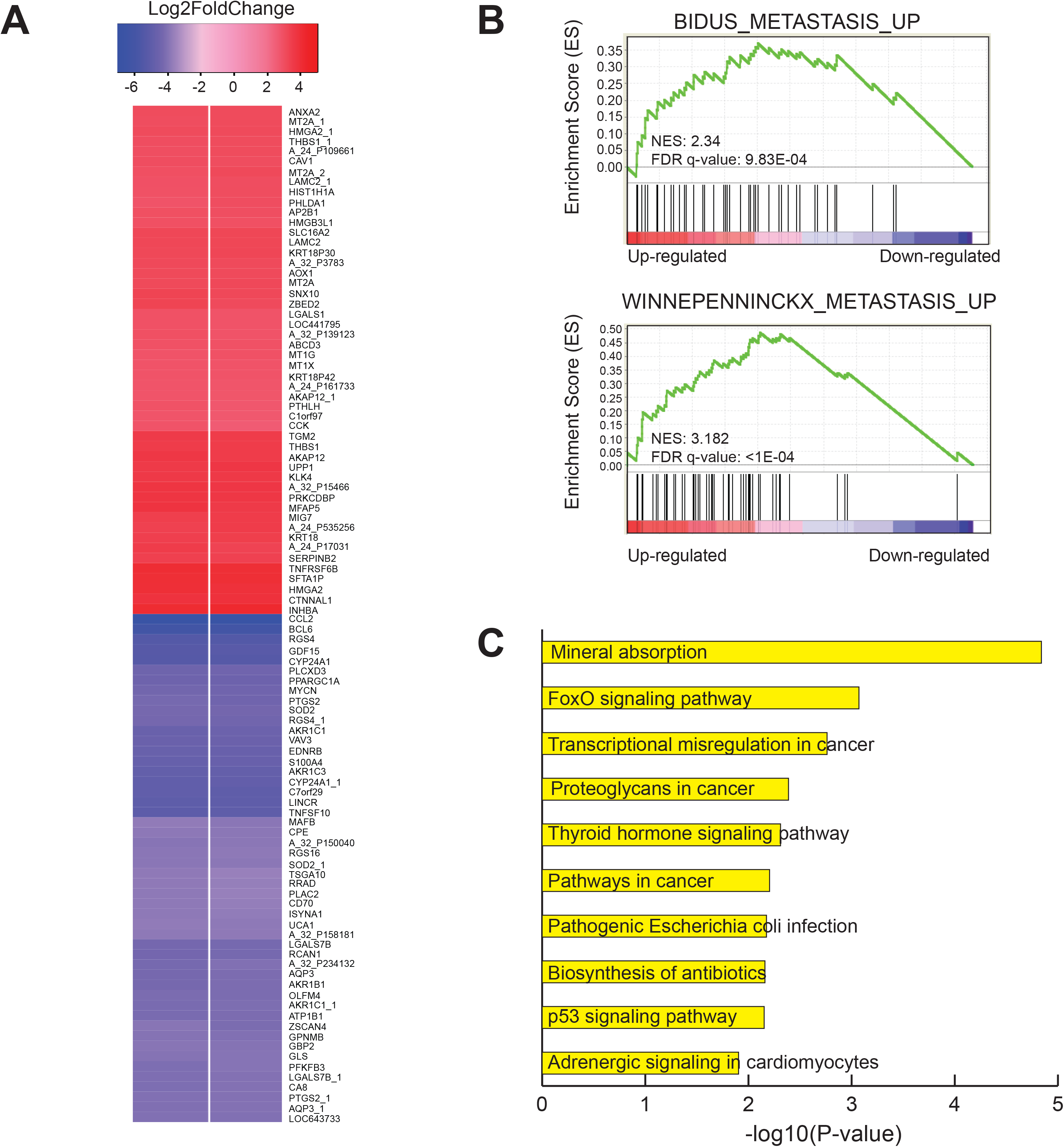
Gene expression analysis of *KLK4-KLKP1*. (A) Heat map showing the top 100 genes differentially expressed in RWPE-1 cells stably transfected with *KLK4-KLKP1* compared to cells transfected with LACZ. The results from 2 independent trials are shown. (B) Gene set enrichment analysis of differentially expressed genes. The genes were enriched in 2 curated gene sets, one involving genes upregulated in endometroid endometrial metastatic tumor “BIDUS_METASTASIS_UP” (top image) and the other including genes overexpressed in melanoma metastatic cancer “WINNEPENNINCKX_METASTASIS_UP” (bottom image). (C) Top 10 KEGG pathways enriched in differentially expressed genes obtained using DAVID tool.

Given the well-established role of androgen receptor (*AR*) in gene expression in prostate cancer [35], we also explored if *AR* is driving the expression of *KLK4-KLKP1* in prostate cancer. Additionally, since we observed concurrent expression of ERG with *KLK4-KLKP1* (Fig. 1G), we also studied if *ERG* is involved in the expression of *KLK4-KLKP1*. Therefore, to identify any AR or ERG binding sequences on *KLK4* or *KLKP1*, we examined data from a previous study where a chromatin immunoprecipitation assay was carried out using antibodies specific to *AR* and *ERG* [36]. Notably, we observed both *AR* and *ERG* binding sites at the fusion junction of *KLKP1* (**Supplementary Fig. S8**), suggesting that both *AR* and *ERG* may modulate the expression of *KLK4-KLKP1* during prostate cancer formation.

For further characterization of the functional role of *KLK4-KLKP1*, we also studied the cellular localization of *KLK4-KLKP1*. We carried out immunofluorescence studies of RWPE-1 cells transfected with adeno-FLAG tagged-*KLK4-KLKP1* using fluorescent anti-FLAG antibody. As a control, cells transfected with adeno-LacZ were used. While cells transfected with adeno-Lacz showed minimal immunofluorescence as expected, notably, we observed colocalization of *KLK4-KLKP1* immunofluorescence signal with 4, 6-diamidino-2-phenylindole (**Supplementary Fig. S9**), indicating that *KLK4-KLKP1* is localized in the nucleus of the cells.

The prostate cancer exclusive expression of *KLK4-KLKP1* in a considerable subset of patients indicates the possible use of *KLK4-KLKP1* as a biomarker for prostate cancer. Therefore, to further explore the potential utility of *KLK4-KLKP1* as a prostate cancer marker, we investigated the association between *KLK4-KLKP1* expression and preoperative PSA of the 659 patients in our cohort. Specifically, we performed a t-test to evaluate difference in log-transformed preoperative PSA between cases with and without *KLK4-KLKP1* expression. Interestingly, patients with *KLK4-KLKP1* expression showed slightly lower preoperative PSA values compared to patients without *KLK4-KLKP1* expression **(Supplementary Fig. S10)**. As a further step, we also analyzed the association between *KLK4-KLKP1* and the time to biochemical recurrence, using multivariable Cox regression model. Patients with *KLK4-KLKP1* showed a lower risk of biochemical recurrence (HR = 0.58; **Supplementary Fig. S11**) after adjusting for age, Gleason grade, and tumor stage. However, the difference in recurrence was not statistically significant (p = 0.12), possibly due to small power as the number of patients showing recurrence was small (n = 49). Additionally, we also analyzed the association of *KLK4-KLKP1* with other clinical and pathological parameters such as family history, tumor stage, tumor volume, metastasis to lymph nodes, perineural invasion and presence of lymph vascular invasion using Pearson’s chi-square test. No statistically significant association was observed between *KLK4-KLKP1* and the clinicopathological variables. Lastly, like TMPRSS2-ERG gene fusions in prostate cancer, we explored the feasibility of detecting *KLK4-KLKP1* in urine samples of prostate cancer patients for noninvasive detection of this marker. We collected urine samples from 90 unselected prostate cancer patients. All patients had confirmed prostate cancer, with most having metastatic or biochemically recurrent disease. Then we screened for *KLK4-KLKP1* transcript using qRT-PCR. As a positive control, RWPE-1 cells stably expressing *KLK4-KLKP1* was used. Importantly, *KLK4-KLKP1* expression was detected in 15 out of 90 (17%) patient samples (**Supplementary Fig. S12**), suggesting the potential for noninvasive detection in patient urine samples. Overall, our study establishes *KLK4-KLKP1* as a recurrent chimeric transcript exclusively expressed in prostate cancer tissues with implications on disease progression and feasibility of being noninvasively detected in patient urine samples.

## DISCUSSION

Given the complex heterogeneous nature of prostate cancer, the identification of distinct patient subgroups based on molecular markers is a necessary step towards targeted disease management. Therefore, in this study we further explored and characterized a pseudogene associated gene fusion *KLK4-KLKP1*. We established that *KLK4-KLKP1* is a recurrent, prostate cancer exclusive fusion transcript that occurs at a significant incidence rate (32%) among prostate cancer patients. Similar to other distinct molecular aberrations such as ETS rearrangements [9] and *SPINK1* mutation [10], *KLK4-KLKP1* was observed only in a subset of prostate cancer patients. However, unlike the mutually exclusive pattern of expression of ETS rearrangements and *SPINK1*, *KLK4-KLKP1* showed concomitant expression with *ERG*, indicating possible cross-talk with ERG. Notably, *KLK4-KLKP1* expression was associated with intact *PTEN* status, suggesting these fusion positive tumors are distinct molecular subtypes from ERG+/PTEN-tumors. Interestingly, full-length normal *KLKP1* transcript showed normal prostate specific expression (GTEX portal) and not in prostate cancer. Furthermore, despite *KLKP1* being categorized as a pseudogene, we showed that *KLK4-KLKP1* is expressed as a full-length protein in a rare phenomenon where gene fusion leads to the inclusion of a pseudogene segment in an expressed protein. Importantly, *KLK4-KLKP1* promoted proliferation, invasion, intravasation and tumor formation, suggesting functional implications on prostate cancer development. Moreover, gene expression studies revealed considerable transcriptional changes in cancer-related genes in cells transfected with *KLK4-KLKP1*, which may indicate that *KLK4-KLKP1* may play a role in transcription during prostate cancer formation. In agreement with a role in transcriptional regulation, *KLK4-KLKP1* was also seen to be localized in the nucleus. Furthermore, both *ERG* and *AR* were found to have binding sites on *KLKP1*, indicating that *KLK4-KLKP1* expression may be *ERG* and *AR* modulated. Finally, we showed that *KLK4-KLKP1* can be easily detected in patient urine samples, suggesting the feasibility for possible future use as a biomarker for early detection of high Gleason grade prostate cancer. Altogether, our study establishes *KLK4-KLKP1* as a novel player in a subset of prostate cancer cases with likely roles in tumor formation.

Long thought to be junk or nonfunctional units of the human genome, pseudogenes have been recently acknowledged to have key cellular roles, particularly in diseases such as cancer [37]. While some pseudogenes are known to be transcribed into non-coding RNA [37], a few pseudogenes have been shown to be even express proteins [20]. Studies have revealed that several different variants of *KLKP1* pseudogene are transcribed exclusively in prostate tissues (**Supplementary Fig. S1**) in an androgen regulated manner [21, 38]. Of the different variants, at least one *KLKP1* variant has been shown to be expressed as a protein in a transfected cell, although not *in vivo* [21]. Even though the variant chimeric transcripts of KLK*4-KLKP1* has been previously described [39, 40], it has not been reported to be expressed as a protein and the functional characteristics have not been validated. Importantly, we verified that *KLK4-KLKP1* is expressed as a full-length protein in both transfected cells and endogenously in castration resistant prostate cancer (PDX), suggesting the occurrence in prostate cancer tissues. In contrast to *KLK4*, which is overexpressed in prostate cancer with roles in cell proliferation, migration and cancer metastasis [41–43], all KLKP1 variants are known to be expressed more in normal prostate tissues compared to prostate cancer [21, 38]. However, *KLK4-KLKP1* is exclusively expressed in prostate cancer with co-occurrence with ERG+ tumors. Thus, our results indicate novel complexity in the *KLK4* and *KLKP1* locus and hint at differential expression of the loci in prostate cancer cells compared to normal prostate cells. Given the presence of *AR* and *ERG* binding sites on *KLKP1* and the previous reports demonstrating *AR* regulation of *KLKP1* expression [21, 38], it is likely that prostate cancer specific expression of *KLK4-KLKP1* is modulated by *AR* and *ERG*. Furthermore, additional variants of *KLK4-KLKP1*, which are different from the *KLK4-KLKP1* transcript observed in prostate cancer, have also been reported in renal cell cancer [40]. While the alternative *KLK4-KLKP1* transcripts were found to occur in a considerable subset of renal cell cancer cases (27%), none of the variants were shown to be expressed as proteins. Thus *KLK4-KLKP1* may be spliced and expressed differently in a tissue specific manner in distinct cancers. Taken together, our results suggest that *KLK4* and *KLKP1* may be a diverse locus that undergo differential splicing and transcription with functional implications in cancer. Consequently, our work highlights unprecedented roles of pseudogenes and complex molecular events involved in cancer.

In agreement with previous reports indicating significant molecular heterogeneity among prostate cancer cases [7], *KLK4-KLKP1* was expressed only in a subset of prostate cancer patients (32%). Additionally, *KLK4-KLKP1* expression was significantly higher in younger patients compared to older prostate cancer patients. Given the oncogenic properties and the transcriptional changes observed with *KLK4-KLKP1*, our results suggest that distinct molecular changes may dictate unique prostate cancer clinical outcomes among patients. Thus, our study further emphasizes the need for subtype specific molecular markers in prostate cancer control. In addition to enhancing cell proliferation, invasion and tumor formation, *KLK4-KLKP1* also caused marked changes in gene expression. Notably, genes affected by *KLK4-KLKP1* were cancer-related and were involved in metastasis of other cancers, implicating a functional role for *KLK4-KLKP1* in prostate cancer. Additionally, ERG was found to have a binding site on *KLK4-KLKP1*. Given that ERG expression was associated with *KLK4-KLKP1*, *ERG* may bind to the *KLKP1* locus and may promote the expression of *KLK4-KLKP1* in a subset of prostate cancer patients.

Even though *KLK4-KLKP1* was implicated in metastatic prostate cancer, the association of *KLK4-KLKP1* with intact *PTEN* status and lower preoperative PSA values suggests indolent disease in prostate cancer patients with *KLK4-KLKP1* expression. However, larger studies exploring the association between *KLK4-KLKP1* expression and prostate cancer clinical outcomes are necessary to establish *KLK4-KLKP1* as a biomarker for prostate cancer. Furthermore, detailed studies are also necessary to fully understand the molecular mechanisms through which *KLK4-KLKP1* promotes prostate cancer formation. Consequently, such studies will explore the potential of *KLK4-KLKP1* as a biomarker and a therapeutic target in prostate cancer, eventually making significant contributions towards achieving effective prostate cancer control.

## MATERIALS AND METHODS

### Tissue microarray construction

Prostatectomy samples collected from 659 patients who underwent radical prostatectomy at Henry Ford Health Systems (HFHS), were reviewed and tissue cores from different regions of the tumor were isolated to construct paraffin embedded tissue microarrays. In most cases, a total of three tissue cores were obtained from each prostatectomy sample. In all cases, appropriate informed consent and Institutional Review Board approval were obtained. The Gleason grade of each tissue core and the race of the patients were reviewed by the study pathologists (NG and SW). Clinical and pathological information of patients such as age, race, family history of prostate cancer, pre-operative PSA, prostatectomy date, Gleason Grade group, tumor stage, cancer status of the lymph nodes, tumor volume, perineural invasion, presence of lymph vascular invasion, last PSA, last PSA date, presence of biochemical recurrence, date of biochemical recurrence were also recorded.

### KLK4-KLKP1 RNA *in situ* Hybridization (RNA-ISH)

RNA-ISH was performed as described previously using RNAscope 2.5 HD Reagent Kit (ACDBio, catalog #322350) according to the manufacturer’s instructions (1). Briefly, after baking, deparaffinization, and target retrieval per manufacturer’s instructions, TMA slides were incubated with target probes for KLK4-KLKP1 (ACDBio, catalog #405501, NM_001136154, region 2933– 3913) for 2 hours at 40°C in a humidity chamber. After detection and color development, slides were washed twice in deionized water and then counterstained in hematoxylin (Agilent DAKO, catalog #K800821-2) for 5 minutes. Slides were washed several times in tap water, then dried, dipped in xylene, and mounted in EcoMount (Fisher, catalog #50–828-32). Next the slides were scanned using a digital imaging system (Aperio Scanner, Leica). The images were reviewed and the RNA-ISH signal on the TMAs was scored. A staining pattern of distinct punctuate cytoplasmic dots was considered as a positive RNA-ISH signal for KLK4-KLKP1 expression. Depending on the intensity of the RNA-ISH staining, a score ranging from +1 to +4 was given to tissue cores with positive RNA-ISH signal, with +1 assigned to the weakest RNA-ISH staining, and +4 given to the cores showing the most intense RNA-ISH staining. A score of 0 was assigned to tissue cores with no visible RNA-ISH staining. The highest score observed among the tissue cores was then assigned to each patient case. If all tissue cores of a patient was 0, the case was recorded as negative.

### Cell culture

HEK-293 cells and prostate benign epithelial cells (RWPE-1, #CRL-11609) were purchased from American Type Culture Collection (Manassas, VA). Primary prostate epithelial cells (PrEC) were purchased from Lonza (Walkersville, MD). HEK-293 cells were cultured in MEM media (Thermo Fisher Scientific, catalog #11095080,) supplemented with 10% FBS (fetal bovine serum, Thermo Fisher Scientific, catalog number #10082147). RWPE-1 cells were cultured in Keratinocyte serum free medium (K-SFM, Gibco™, Thermo Fisher Scientific, catalog #17005-042, Carlsbad, CA) supplemented with Bovine Pituitary Extract (BPE, 0.05 mg/ml, Thermo Fisher Scientific, catalog #17005-042), human recombinant Epidermal Growth Factor 1-53 (EGF 1-53, 5 ng/ml, Thermo Fisher Scientific, catalog #17005-042) and 1% penicillin/streptomycin. PrEC cells were cultured in Prostate Epithelial Cell Basal Medium (PrEGM) supplemented with Prostate Epithelial Cell Growth Kit (Clonetics™ PrEGM™, BulletKit™, Lonza). All cell cultures were maintained at 37°C in an incubator with a controlled humidified atmosphere composed of 95% air and 5% CO2.

### *In vitro* overexpression of KLK4-KLKP1

KLK4-KLKP1 cDNA was PCR amplified using a forward primer with DDK tag and a reverse primer from KLK4-KLKP1 template and was cloned into Gateway expression system (Life Technologies). To generate lentiviral and adenoviral constructs, PCR8-KLK4-KLKP1 (DDK tagged) was recombined with pLenti6/V5-Dest™ (Life Technologies) or pAD/CMV/V5-Dest™ (Life Technologies), respectively using LR Clonase II (Life Technologies). For transient overexpression in HEK-293, RWPE-1 and PrEC cells, adenoviruses carrying KLK4-KLKP1, EZH2 or lacZ were added to the culture media after cells reached 50-70% confluency. At the same time, cells were treated with or without bortezomib (100nM in ethanol, 10 µL, Cayman Chemical, catalog #10008822). After incubation for 48 hours at 370C, cells were harvested by scraping. For stable overexpression, RWPE-1 cells were infected with lentiviruses expressing KLK4-KLKP1 or lacZ, and stable clones were selected with blasticidin (3.5 μg/ml, Sigma-aldrich, MO, USA). Lenti and adeno viruses were generated by the University of Michigan Vector Core (Ann Arbor, MI, USA).

### Western blotting

Harvested cells were spun down (1000 rpm, 5 min, 4 °C). For HEK-293 cells, the cell pellet was re-suspended in RIPA lysis buffer (Thermo Fisher Scientific, catalog #PI89900) supplemented with protease inhibitor (1X, genDEPOT, catalog #50-101-5488). For RWPE-1 cells, NP-40 lysis buffer (Boston BioProducts, Ashland, MA) with protease inhibitor was used to lyse the cells. With xenograft tissues, frozen tissues were cut into small pieces and then sonicated on ice in RIPA lysis buffer. The debris from cells or tissues were removed by centrifugation (13.2 rpm, 10 minutes, 4 0C). Protein concentration of the supernatant was determined using Micro BCA protein assay kit (Thermo Fisher Scientific, catalog #23235). The lysates were separated on a 12% SDS-PAGE or a NuPAGE™ 4-12% Bis-Tris protein gel. After separation, proteins were transferred onto a PVDF membrane (Milipore Immobilon-P, Fisher, catalog #IPVH00010). Then the membranes were probed with specific antibodies: Flag (Sigma, catalog #F1804), KLK4/KLKP1 (Eurogentec custom synthesized antibody) and β-actin (Sigma, catalog #A2228). The membranes were visualized on an imaging system (ChemiDoc, BIO-RAD) using a chemiluminescence developing kit (Clarity™ Western ECL Blotting Substrates, BIO-RAD, catalog #1705060).

### Measurement of cell proliferation

Cell proliferation was measured by cell counting. For this, stable RWPE-1 cells overexpressing KLK4-KLKP1 (DDK-tagged) or lacZ were used. The cells were seeded at a density of 10 000 cells per well in 24-well plates (n=3). Next, the cells were trypsinized and counted at specified time points by Z2 Coulter particle counter (Beckman Coulter, Brea, CA, USA). LacZ cells were served as controls. Each experiment has been performed with three replicates per sample.

### Matrigel invasion assay

Matrigel invasion assays were performed using BD BioCoat Matrigel matrix (Corning Life Sciences, Tewksbury, MA, USA). The parental and transfected clones of RWPE-1 and PrEC cells were seeded at 1 × 105 cells in serum-free medium in the upper chamber of a 24-well culture plate. The lower chamber containing respective medium was supplemented with 10% serum as a chemoattractant. After 48 h, the non-invading cells and Matrigel matrix from the upper side of the chamber were gently wiped with a cotton swab. Invasive cells located on the lower side of the chamber were stained with 0.2% crystal violet in methanol, air-dried and photographed using an inverted microscope (4x). Invasion was quantified by colorimetric assay or by counting the number of cells. For colorimetric assays, the inserts were treated with 150 μl of 10% acetic acid and the absorbance measured at 560 nm.

### Chicken Chorioallantoic Membrane Assay (CAM) assay

CAM assay was performed as described earlier [31]. Briefly, fertilized eggs were incubated in a rotary humidified incubator at 38°C for 10 days. CAM was dropped by making two holes, one through the eggshell into the air sac and a second hole near the allantoic vein that penetrates the eggshell membrane but not the CAM. Subsequently a cutoff wheel (Dremel) was used to cut a 1 cm2 window to expose the underlying CAM near the allantoic vein. After 3 days of implanting the 2*10^6^ cells in 50 ul medium on the top of each egg, lower CAM was harvested and analyzed for the presence of tumor cells by quantitative human Alu-specific PCR. Genomic DNA from lower CAM and livers were prepared using Puregene DNA purification system (Qiagen USA) and quantification of human-Alu was performed as described (Ref). After 7 days of implantation, extraembryonic tumors were isolated and weighed. An average of 8 eggs per group was used in ea

### Gene expression microarray analysis

Two-channel microarray experiment was performed with two replicates using the Agilent Whole Human Genome Oligo Microarray (Agilent, catalog #G4851C Whole Human Genome Microarray 8×60K). Raw data from each replicates were independently processed using Bioconductor packages. “agilp” Bioconductor package (1) was used to apply loess normalization on raw expression values. Fold change for each probe was obtained by taking difference of loess-normalized, log-2–transformed signal intensity between sample with KLK4-KLKP1 gene fusion and control sample. Probes showing differential expression in both two-channel experiments were considered for functional analysis. In total, 1956 probes were up-regulated (with Log2FC >=1) and 1918 probes were down-regulated (with log2FC <= −1) in KLK4-KLKP1 gene fusion sample. Heatmap of differentially expressed genes was created using heatmap.2 of “gplots” R package.

### Gene set enrichment analysis (GSEA)

Gene set enrichment analysis (GSEA) was performed using the curated gene sets [C2] (n=1267) from Molecular Signature Database (MSigDB v5.0) provided by Broad institute (2) Differentially expressed genes were ranked by average log2FC from two arrays and submitted to GSEAPreranked module in GSEA software.

### KEGG pathway analysis

DAVID (Database for Annotation, Visualization and Integrated Discovery) v6.8 (3) was used to identify enriched KEGG pathways in these differentially expressed genes. With default parameters (gene count of 2 and EASE of 0.1), functional annotation chart was obtained and KEGG pathways with p-value <0.05 were considered to be enriched.

### Screening of KLK4-KLKP1 in the urine samples of prostate cancer patients

Random urine samples were collected with informed consent and Institutional Review Board approval from PCa patients visiting the Hematology Oncology clinic at Henry Ford hospital in Detroit, MI.RNA was isolated using ZR urine RNA isolation kitTM (Zymo Research, catalog # R1038 & R1039) according to manufacturer’s instructions. cDNA synthesis and qRT-PCR were performed as described earlier.

### Statistical analysis

Pearson’s chi-square test was used to evaluate the association of KLK4-KLKP1 fusion with race, age, Gleason score and other molecular markers. For association between KLK4-KLKP1 and pre-operative PSA, two-sample t-test was performed to evaluate difference in log-transformed pre-operative PSA between KLK4-KLKP1 positive and negative cases. Multivariable Cox regression was used to estimate the association between KLK4-KLKP1 and the risk of biochemical recurrence. Cox regression model was adjusted for patients’ age group (<50; >=50), Gleason score (6 or 3+4; 4+3 or 8+), and tumor stage (pT2; pT3 or pT4). For all analyses, a p-value of <.05 was considered statistically significant. All analyses were performed using the Statistical Analysis System (SAS) statistical software package, version 9.1.3. For the rest of the experiments, Student’s two-sample t-test was used to determine significant differences between two groups. P-values <0.05 were considered significant.

## ACKNOWLEDGEMENTS

We would like to thank Mireya Diaz-Insua for helping to identify the prostate patient cohort. Natalia Draga and Jingli Yang for making the prostate tissue microarray. We thank the University of Michigan Sequencing Core and Vector Core facilities for their assistance in sequencing of the clones and construction of the adenoviral and lentiviral constructs used in this study. Arul Chinnaiyan for his support at the Michigan Center for Translational Pathology, University of Michigan during the early phase of the project.

## AUTHOR CONTRIBUTIONS

Conception and design: NP

Development and methodology: SV, NP

Acquisition of data: BVSKC, PDA, SC

Analysis and interpretation of data: SK, JL, KHHW, DSC, NP, SV, PDA, NG, SW, DC, NP

Writing, review and/or revision of the manuscript: NP, PDA

Administrative, technical, or material support: NN, JP, HS, CR, MM,

Study supervision: SV, NP

Other:

## Notes

Conflicts of Interest: None

## REFERENCES

[1] Cronin KA, Lake AJ, Scott S, Sherman RL, Noone AM, Howlader N, Henley SJ, Anderson RN, Firth AU, Ma J, et al. (2018). Annual Report to the Nation on the Status of Cancer, part I: National cancer statistics Cancer 124, 2785–2800.

[2] Bray F, Ferlay J, Soerjomataram I, Siegel RL, Torre LA, Jemal A (2018). Global cancer statistics 2018: GLOBOCAN estimates of incidence and mortality worldwide for 36 cancers in 185 countries CA Cancer J Clin 68, 394–424.

[3] Abate-Shen C, Shen MM (2000). Molecular genetics of prostate cancer Genes & development 14, 2410–2434.

[4] Arora R, Koch MO, Eble JN, Ulbright TM, Li L, Cheng L (2004). Heterogeneity of Gleason grade in multifocal adenocarcinoma of the prostate Cancer 100, 2362–2366.

[5] Cheng L, MacLennan GT, Lopez-Beltran A, Montironi R (2012). Anatomic, morphologic and genetic heterogeneity of prostate cancer: implications for clinical practice Expert Rev Anticancer Ther 12, 1371–1374.

[6] Cancer Genome Atlas Research N (2015). The molecular taxonomy of primary prostate cancer Cell 163, 1011–1025.

[7] Tomlins SA, Alshalalfa M, Davicioni E, Erho N, Yousefi K, Zhao S, Haddad Z, Den RB, Dicker AP, Trock BJ, et al. (2015). Characterization of 1577 primary prostate cancers reveals novel biological and clinicopathologic insights into molecular subtypes Eur Urol 68, 555–567.

[8] Yang L, Wang S, Zhou M, Chen X, Jiang W, Zuo Y, Lv Y (2017). Molecular classification of prostate adenocarcinoma by the integrated somatic mutation profiles and molecular network Sci Rep 7, 738.

[9] Tomlins SA, Bjartell A, Chinnaiyan AM, Jenster G, Nam RK, Rubin MA, Schalken JA (2009). ETS gene fusions in prostate cancer: from discovery to daily clinical practice Eur Urol 56, 275–286.

[10] Tomlins SA, Rhodes DR, Yu J, Varambally S, Mehra R, Perner S, Demichelis F, Helgeson BE, Laxman B, Morris DS, et al. (2008). The role of SPINK1 in ETS rearrangement-negative prostate cancers Cancer cell 13, 519–528.

[11] Palanisamy N, Ateeq B, Kalyana-Sundaram S, Pflueger D, Ramnarayanan K, Shankar S, Han B, Cao Q, Cao X, Suleman K, et al. (2010). Rearrangements of the RAF kinase pathway in prostate cancer, gastric cancer and melanoma Nat Med 16, 793–798.

[12] Baena E, Shao Z, Linn DE, Glass K, Hamblen MJ, Fujiwara Y, Kim J, Nguyen M, Zhang X, Godinho FJ, et al. (2013). ETV1 directs androgen metabolism and confers aggressive prostate cancer in targeted mice and patients Genes Dev 27, 683–698.

[13] Nam RK, Sugar L, Yang W, Srivastava S, Klotz LH, Yang LY, Stanimirovic A, Encioiu E, Neill M, Loblaw DA, et al. (2007). Expression of the TMPRSS2:ERG fusion gene predicts cancer recurrence after surgery for localised prostate cancer Br J Cancer 97, 1690–1695.

[14] Gleason DF (1992). Histologic grading of prostate cancer: a perspective Hum Pathol 23, 273–279.

[15] Adhyam M, Gupta AK (2012). A Review on the Clinical Utility of PSA in Cancer Prostate Indian J Surg Oncol 3, 120–129.

[16] Carter HB, Partin AW, Walsh PC, Trock BJ, Veltri RW, Nelson WG, Coffey DS, Singer EA, Epstein JI (2012). Gleason score 6 adenocarcinoma: should it be labeled as cancer? J Clin Oncol 30, 4294–4296.

[17] Tan SH, Petrovics G, Srivastava S (2018). Prostate cancer genomics: recent advances and the prevailing underrepresentation from racial and ethnic minorities Int J Mol Sci 19, E1255.

[18] Kalyana-Sundaram S, Kumar-Sinha C, Shankar S, Robinson DR, Wu YM, Cao X, Asangani IA, Kothari V, Prensner JR, Lonigro RJ, et al. (2012). Expressed pseudogenes in the transcriptional landscape of human cancers Cell 149, 1622–1634.

[19] Poliseno L (2012). Pseudogenes: newly discovered players in human cancer Sci Signal 5, re5.

[20] Brosch M, Saunders GI, Frankish A, Collins MO, Yu L, Wright J, Verstraten R, Adams DJ, Harrow J, Choudhary JS, et al. (2011). Shotgun proteomics aids discovery of novel protein-coding genes, alternative splicing, and “resurrected” pseudogenes in the mouse genome Genome Res 21, 756–767.

[21] Kaushal A, Myers SA, Dong Y, Lai J, Tan OL, Bui LT, Hunt ML, Digby MR, Samaratunga H, Gardiner RA, et al. (2008). A novel transcript from the KLKP1 gene is androgen regulated, down-regulated during prostate cancer progression and encodes the first non-serine protease identified from the human kallikrein gene locus Prostate 68, 381–399.

[22] Clements J, Hooper J, Dong Y, Harvey T (2001). The expanded human kallikrein (KLK) gene family: genomic organisation, tissue-specific expression and potential functions Biol Chem 382, 5–14.

[23] Warrick JI, Tomlins SA, Carskadon SL, Young AM, Siddiqui J, Wei JT, Chinnaiyan AM, Kunju LP, Palanisamy N (2014). Evaluation of tissue PCA3 expression in prostate cancer by RNA in situ hybridization--a correlative study with urine PCA3 and TMPRSS2-ERG Mod Pathol 27, 609–620.

[24] Leinonen KA, Saramaki OR, Furusato B, Kimura T, Takahashi H, Egawa S, Suzuki H, Keiger K, Ho Hahm S, Isaacs WB, et al. (2013). Loss of PTEN is associated with aggressive behavior in ERG-positive prostate cancer Cancer Epidemiol Biomarkers Prev 22, 2333–2344.

[25] Ahearn TU, Pettersson A, Ebot EM, Gerke T, Graff RE, Morais CL, Hicks JL, Wilson KM, Rider JR, Sesso HD, et al. (2016). A prospective investigation of PTEN loss and ERG expression in lethal prostate cancer J Natl Cancer Inst 108, djv346.

[26] Fontugne J, Lee D, Cantaloni C, Barbieri CE, Caffo O, Hanspeter E, Mazzoleni G, Dalla Palma P, Rubin MA, Fellin G, et al. (2014). Recurrent prostate cancer genomic alterations predict response to brachytherapy treatment Cancer Epidemiol Biomarkers Prev 23, 594–600.

[27] Bismar TA, Hegazy S, Feng Z, Yu D, Donnelly B, Palanisamy N, Trock BJ (2018). Clinical utility of assessing PTEN and ERG protein expression in prostate cancer patients: a proposed method for risk stratification J Cancer Res Clin Oncol 144, 2117–2125.

[28] Navone NM, van Weerden WM, Vessella RL, Williams ED, Wang Y, Isaacs JT, Nguyen HM, Culig Z, van der Pluijm G, Rentsch CA, et al. (2018). Movember GAP1 PDX project: An international collection of serially transplantable prostate cancer patient-derived xenograft (PDX) models Prostate 78, 1262–1282.

[29] Ren G, Baritaki S, Marathe H, Feng J, Park S, Beach S, Bazeley PS, Beshir AB, Fenteany G, Mehra R, et al. (2012). Polycomb protein EZH2 regulates tumor invasion via the transcriptional repression of the metastasis suppressor RKIP in breast and prostate cancer Cancer Res 72, 3091–3104.

[30] Bryant RJ, Cross NA, Eaton CL, Hamdy FC, Cunliffe VT (2007). EZH2 promotes proliferation and invasiveness of prostate cancer cells Prostate 67, 547–556.

[31] Chakravarthi BV, Goswami MT, Pathi SS, Robinson AD, Cieslik M, Chandrashekar DS, Agarwal S, Siddiqui J, Daignault S, Carskadon SL, et al. (2016). MicroRNA-101 regulated transcriptional modulator SUB1 plays a role in prostate cancer Oncogene 35, 6330–6340.

[32] Chakravarthi BV, Pathi SS, Goswami MT, Cieslik M, Zheng H, Nallasivam S, Arekapudi SR, Jing X, Siddiqui J, Athanikar J, et al. (2014). The miR-124-prolyl hydroxylase P4HA1-MMP1 axis plays a critical role in prostate cancer progression Oncotarget 5, 6654–6669.

[33] Subramanian A, Tamayo P, Mootha VK, Mukherjee S, Ebert BL, Gillette MA, Paulovich A, Pomeroy SL, Golub TR, Lander ES, et al. (2005). Gene set enrichment analysis: a knowledge-based approach for interpreting genome-wide expression profiles Proc Natl Acad Sci U S A 102, 15545–15550.

[34] Huang DW, Sherman BT, Tan Q, Kir J, Liu D, Bryant D, Guo Y, Stephens R, Baseler MW, Lane HC, et al. (2007). DAVID Bioinformatics Resources: expanded annotation database and novel algorithms to better extract biology from large gene lists Nucleic Acids Res 35, W169–175.

[35] Heinlein CA, Chang C (2004). Androgen receptor in prostate cancer Endocr Rev 25, 276–308.

[36] Yu J, Yu J, Mani RS, Cao Q, Brenner CJ, Cao X, Wang X, Wu L, Li J, Hu M, et al. (2010). An integrated network of androgen receptor, polycomb, and TMPRSS2-ERG gene fusions in prostate cancer progression Cancer Cell 17, 443–454.

[37] Pink RC, Wicks K, Caley DP, Punch EK, Jacobs L, Carter DR (2011). Pseudogenes: pseudo-functional or key regulators in health and disease? RNA 17, 792–798.

[38] Lu W, Zhou D, Glusman G, Utleg AG, White JT, Nelson PS, Vasicek TJ, Hood L, Lin B (2006). KLK31P is a novel androgen regulated and transcribed pseudogene of kallikreins that is expressed at lower levels in prostate cancer cells than in normal prostate cells Prostate 66, 936–944.

[39] Lai J, Lehman ML, Dinger ME, Hendy SC, Mercer TR, Seim I, Lawrence MG, Mattick JS, Clements JA, Nelson CC (2010). A variant of the KLK4 gene is expressed as a cis sense-antisense chimeric transcript in prostate cancer cells RNA 16, 1156–1166.

[40] Pflueger D, Mittmann C, Dehler S, Rubin MA, Moch H, Schraml P (2015). Functional characterization of BC039389-GATM and KLK4-KRSP1 chimeric read-through transcripts which are up-regulated in renal cell cancer BMC Genomics 16, 247.

[41] Veveris-Lowe TL, Lawrence MG, Collard RL, Bui L, Herington AC, Nicol DL, Clements JA (2005). Kallikrein 4 (hK4) and prostate-specific antigen (PSA) are associated with the loss of E-cadherin and an epithelial-mesenchymal transition (EMT)-like effect in prostate cancer cells Endocr Relat Cancer 12, 631–643.

[42] Klokk TI, Kilander A, Xi Z, Waehre H, Risberg B, Danielsen HE, Saatcioglu F (2007). Kallikrein 4 is a proliferative factor that is overexpressed in prostate cancer Cancer Res 67, 5221–5230.

[43] Gao J, Collard RL, Bui L, Herington AC, Nicol DL, Clements JA (2007). Kallikrein 4 is a potential mediator of cellular interactions between cancer cells and osteoblasts in metastatic prostate cancer Prostate 67, 348–360.

