## Supplementary material for "Pseudogene associated recurrent gene fusion in prostate cancer": Suppl figure 1-12

**Figure S1**

**A**

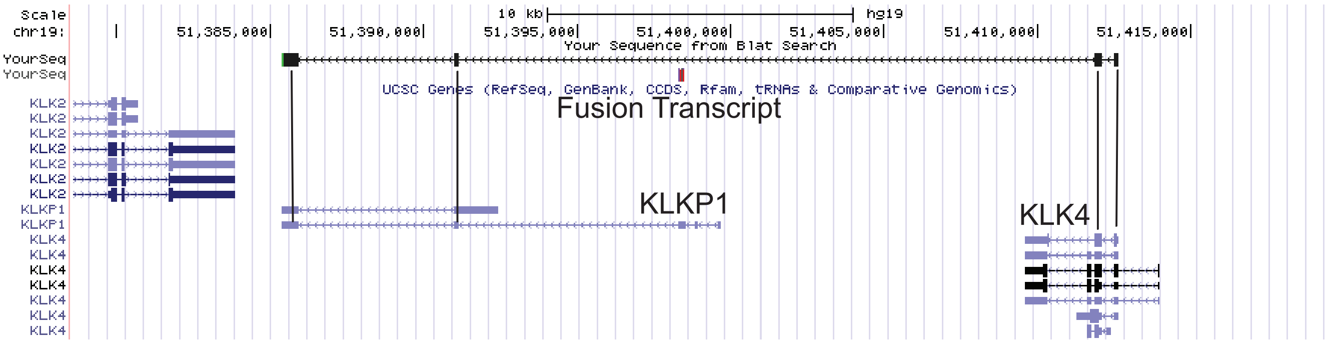


**B**

gatcgctcgtctctggtagctgcagccaaatcataaacggcgaggactgcagcccgcactcgcagccctggcaggcggcactggtc*atggaa
aacgaattgttctgctcgggcgtcctggtgcatccgcagtgggtgctgtcagccgcacactgtttccagaactcctacaccatcgggctgggcctgc
acagtcttgaggccgaccaagagccagggagccagatggtggaggccagcctctccgtacggcacccagagtacaacagacccttgctcgct
aacgacctcatgctcatcaagttggacgaatccgtgtccgagtctgacaccatccggagcatcagcattgcttcgcagtgccctaccgcggggaa
ctcttgcctcgtttctggctggggtctgctggcgaacgATGCTGTGATTGCCATCCAGTCCCAGACTGTGGGAGGCTGG
GAGTGTGAGAAGCTTTCCCAACCCTGGCAGGGTTGTACCATTTCGGCAACTTCCAGTGCAAGGACG
TCCTGCTGCATCCTCACTGGGTGCTCACTACTGCTCACTGCATCACCCGGAACACTGTGA*TCAACT
AGCCAGCACCATAGTTCTCCGAAGTCAGACTATCATGATTACTGTGTTGACTGTGCTGTCTATTGTAC
TAACCATGCCGATGTTTAGGTGAAATTAGCGTCACTTGGCCTCAACCATCTTGGTATCCAGTTATCCT
CACTGAATTGAGATTTCCTGCTTCAGTGTCAGCCATTCCCACATAATTTCTGACCTACAGAGGTGAGG
GATCATATAGCTCTTCAAGGATGCTGGTACTCCCCTCACAAATTCATTTCTCCTGTTGTAGTGAAAGGT
GCGCCCTCTGGAGCCTCCCAGGGTGGGTGTGCAGGTCACAATGATGAATGTATGATCGTGTTCCCAT
TACCCAAAGCCTTTAAATCCCTCATGCTCAGTACACCAGGGCAGGTCTAGCATTTCTTCATTTAGTGTA
TGCTGTCCATTCATGCAACCACCTCAGGACTCCTGGATTCTCTGCCTAGTTGAGCTCCTGCATGCTG
CCTCCTTGGGGAGGTGAGGGAGAGGGCCCATGGTTCAATGGGATCTGTGCAGTTGTAACACATTAG
GTGCTTAATAAACAGAAGCTGTGATGTTAAaaaaaaaaaaaaaaaaaaaaa

**C**

1 ATG GAA AAC GAA TTG TTC TGC TCG GGC GTC CTG GTG CAT CCG CAG 45
**M E N E L F C S G V L V H P Q**
46 TGG GTG CTG TCA GCC GCA CAC TGT TTC CAG AAC TCC TAC ACC ATC 90
**W V L S A A H C F Q N S Y T I**
91 GGG CTG GGC CTG CAC AGT CTT GAG GCC GAC CAA GAG CCA GGG AGC 135
**G L G L H S L E A D Q E P G S**
136 CAG ATG GTG GAG GCC AGC CTC TCC GTA CGG CAC CCA GAG TAC AAC 180
**Q M V E A S L S V R H P E Y N**
181 AGA CCC TTG CTC GCT AAC GAC CTC ATG CTC ATC AAG TTG GAC GAA 225
**R P L L A N D L M L I K L D E**
226 TCC GTG TCC GAG TCT GAC ACC ATC CGG AGC ATC AGC ATT GCT TCG 270
**S V S E S D T I R S I S I A S**
271 CAG TGC CCT ACC GCG GGG AAC TCT TGC CTC GTT TCT GGC TGG GGT 315
**Q C P T A G N S C L V S G W G**
316 CTG CTG GCG AAC GAT GCT GTG ATT GCC ATC CAG TCC CAG ACT GTG 360
**L L A N D A V I A I Q S Q T V**
361 GGA GGC TGG GAG TGT GAG AAG CTT TCC CAA CCC TGG CAG GGT TGT 405
**G G W E C E K L S Q P W Q G C**
406 ACC ATT TCG GCA ACT TCC AGT GCA AGG ACG TCC TGC TGC ATC CTC 450
**T I S A T S S A R T S C C I L**
451 ACT GGG TGC TCA CTA CTG CTC ACT GCA TCA CCC GGA ACA CTG TGA 495
**T G C S L L L T A S P G T L ***

**Figure S1:** Genomic organization of KLK4-KLKP1 fusion transcript. A) Genomic alignment of the cloned fusion transcript in UCSC genome browser (GRCh37/hg19). Sequence alignment corresponded to the first two exons of KLK4 and exons 4-5 of KLKP1. B) Nucleotide sequence of the fusion transcript. KLK4 sequence is shown in red letters and KLKP1 sequence is shown in blue letters. The sequence shown within the* correspond to the open reading frame (ORF). C) Translation of the open reading frame sequence (495bp) into amino acid sequence (164aa). Aminoacids shown in red are derived from the KLKP1 sequence (55aa).

### Figure S2

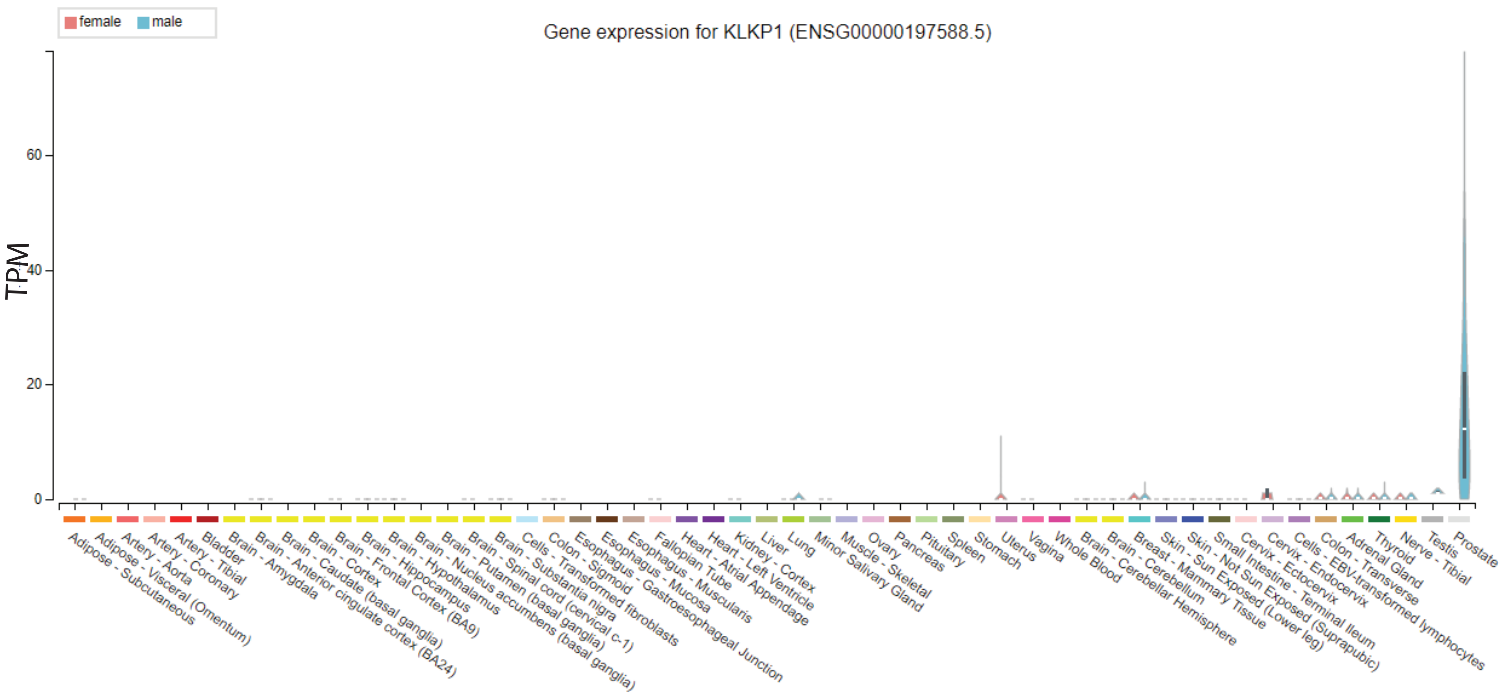


**Figure S2:** The exclusive expression of full length KLKP1 in normal prostate tissue - GTEx portal (https://gtexportal.org/home/gene/KLKP1).

#### Figure S3

A

B

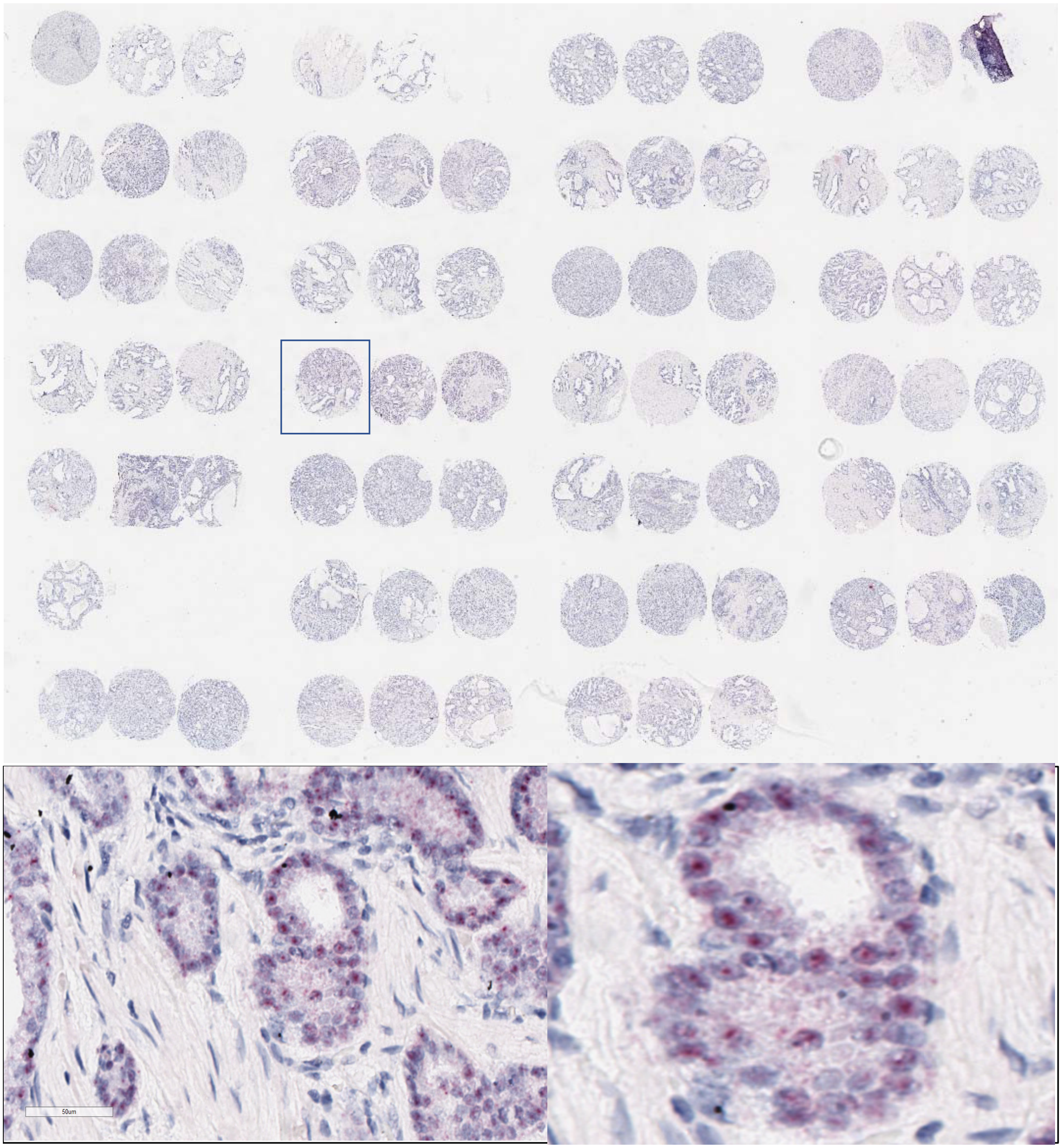


**Figure S3:** A) Representative image of a prostate tissue microarray (TMA). Each patient is represented with triplicate cores. TMA’s are stained with RNA ISH probe for KLK4-KLKP1. B) Enlarge view of a tissue core positive for KLK4-KLKP1 expression. RNA ISH signals appear as distinct punctate dots for each mRNA transcript. Positive signals are specific to the tumor area and the stromal areas do not show any signal indicating the specificity of the RNA ISH probe.

**Figure S4**

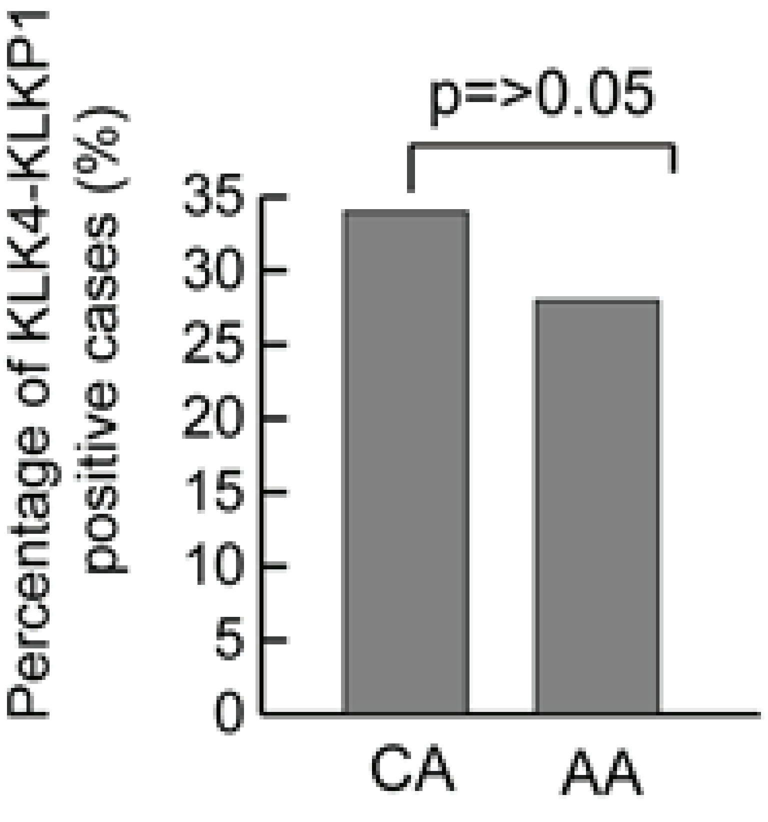


**Figure S4:** KLK4-KLKP1 is expressed equally among CA and AA patients. The percentage of cases with positive KLK4-KLKP1 RNA-ISH signal in CA and AA groups is shown. P Value was calculated using Pearson's chi-square test.

### Figure S5

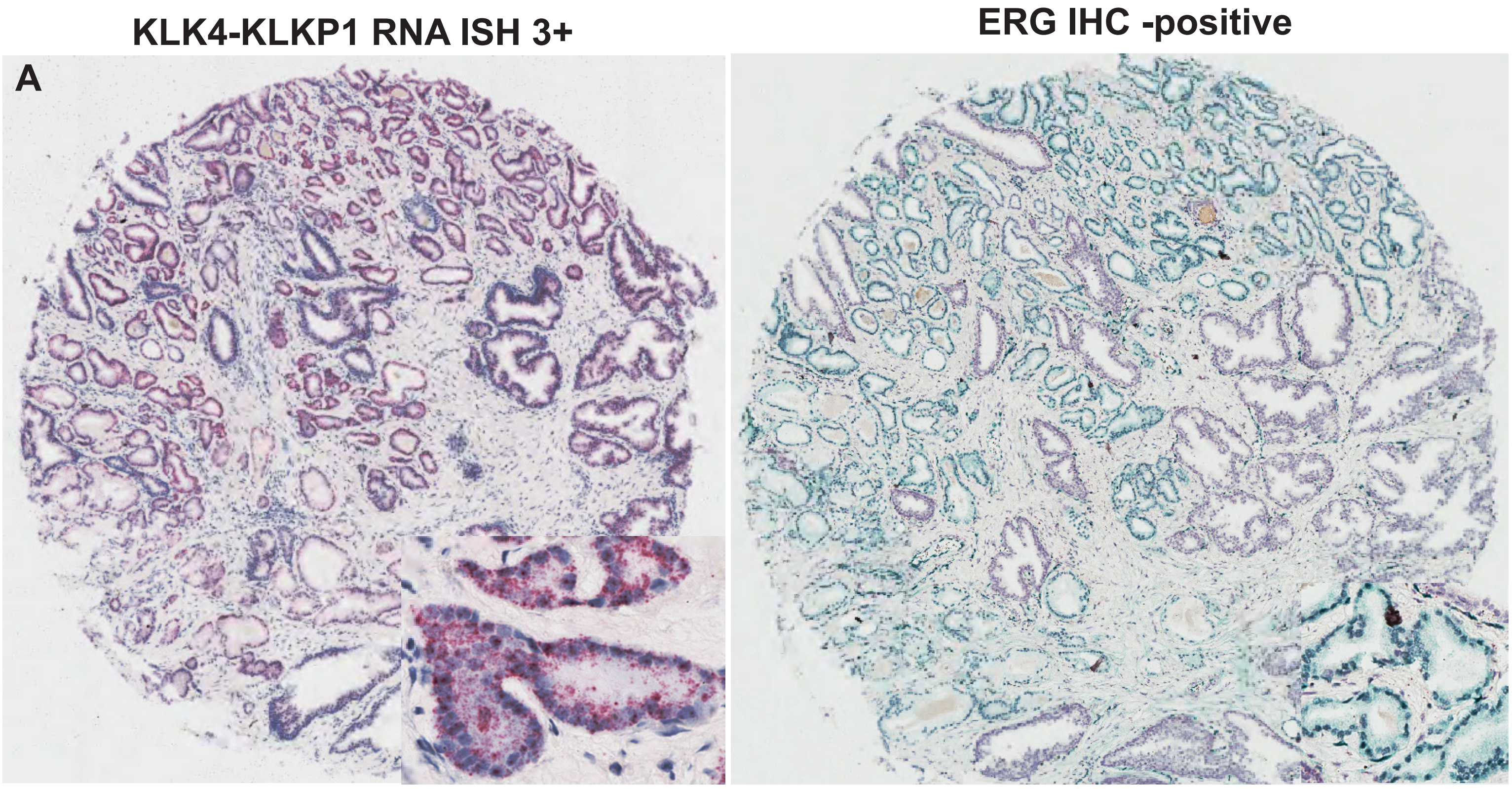


**Figure S5**: Concomitant expression of KLK4-KLKP1 and ERG. A) RNA ISH showing tumor specific expression of KLK4-KLKP1. B) On a consecutive slide from the same patient stained with ERG immunohistochemistry revealed tumor specific staining of the same tumor foci positive for KLK4-KLKP1. Insets showing enlarged view of a tumor area positive for both markers.

### Figure S6

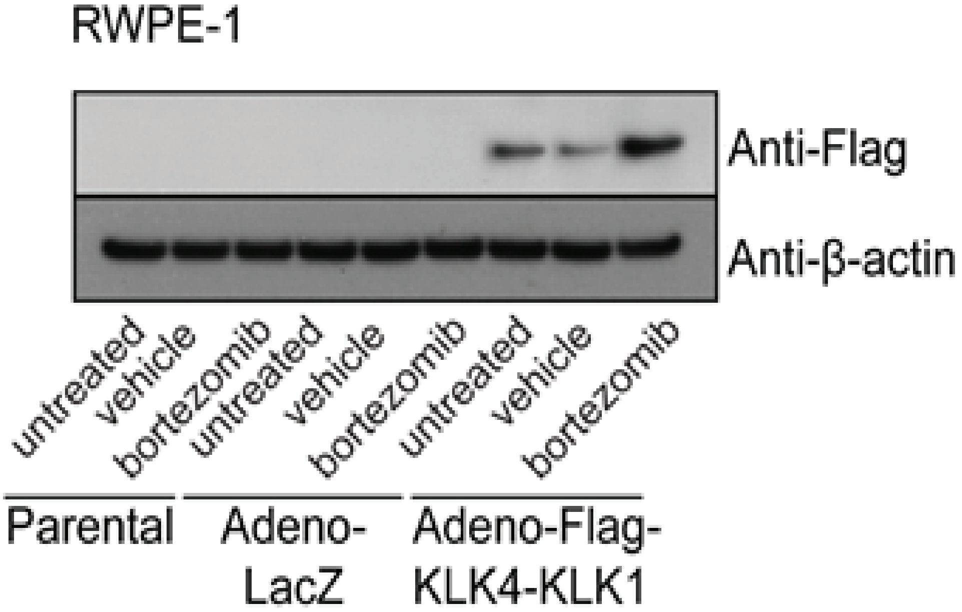


**Figure S6:** Western blot analysis of KLK4-KLKP1 in RWPE-1 cells. Either untreated RWPE-1 cells or cells treated with bortezomib or vehicle (ethanol) were transfected with adeno viral vector carrying flag tagged-KLK4-KLKP1 (Adeno-Flag-KLK4-KLKP1). As controls, cells transfected with adeno viral vector carrying LacZ (Adeno-LacZ) and untransfected cells (Parental) were also used. Cells were lysed, proteins were separated on SDS-PAGE and visualized using anti-Flag and anti-β-actin antibodies after transfer onto a PVDF membrane

### Figure S7

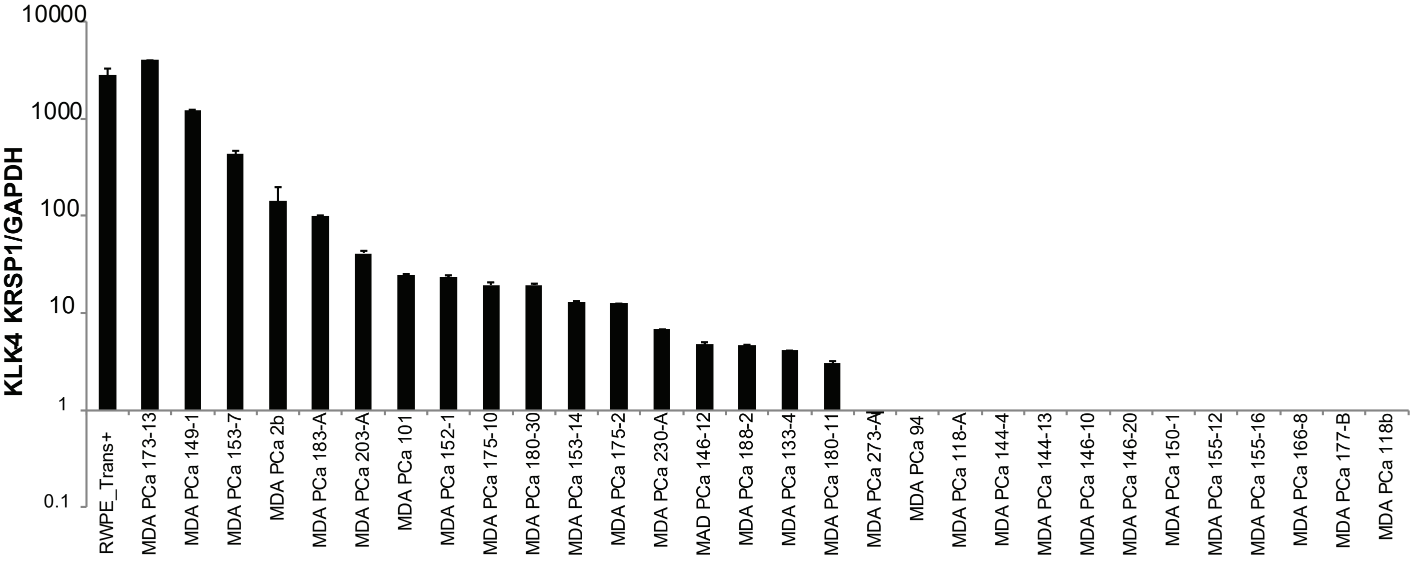


**Figure S7**: Screening of Prostate cancer patient derived xenografts for KLK4-KLKP1 by qRT-PCR. RWPE cells transfected with KLK4-KLKP1 construct was used as a positive control. The relative expression was calculated against GAPDH level.

### Figure S8

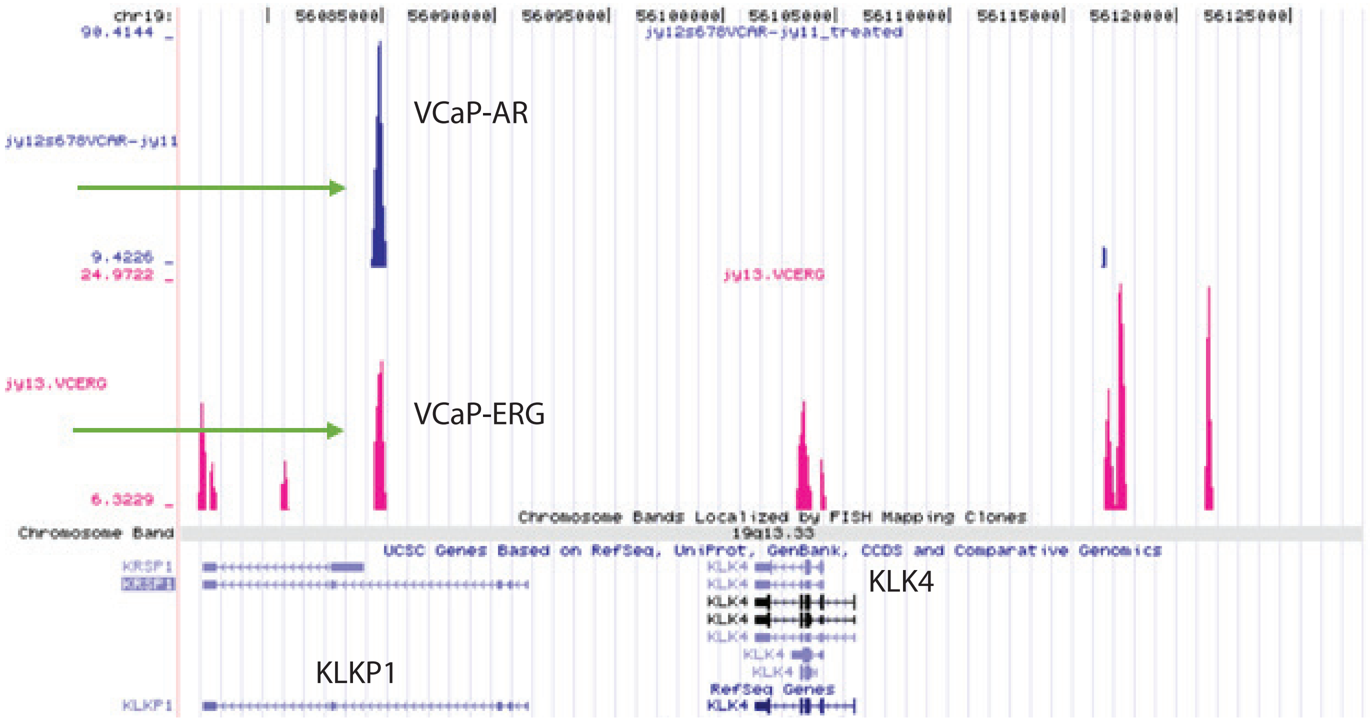


**Figure S8:** The representative image of ERG and AR binding in the KLKP1 region. AR binding regions (blue) and ERG binding regions (pink) in the KLKP1 and KLK4 locus are shown on the UCSC genome browser. The green arrows indicate the AR and ERG binding regions on KLKP1.

### Figure 9

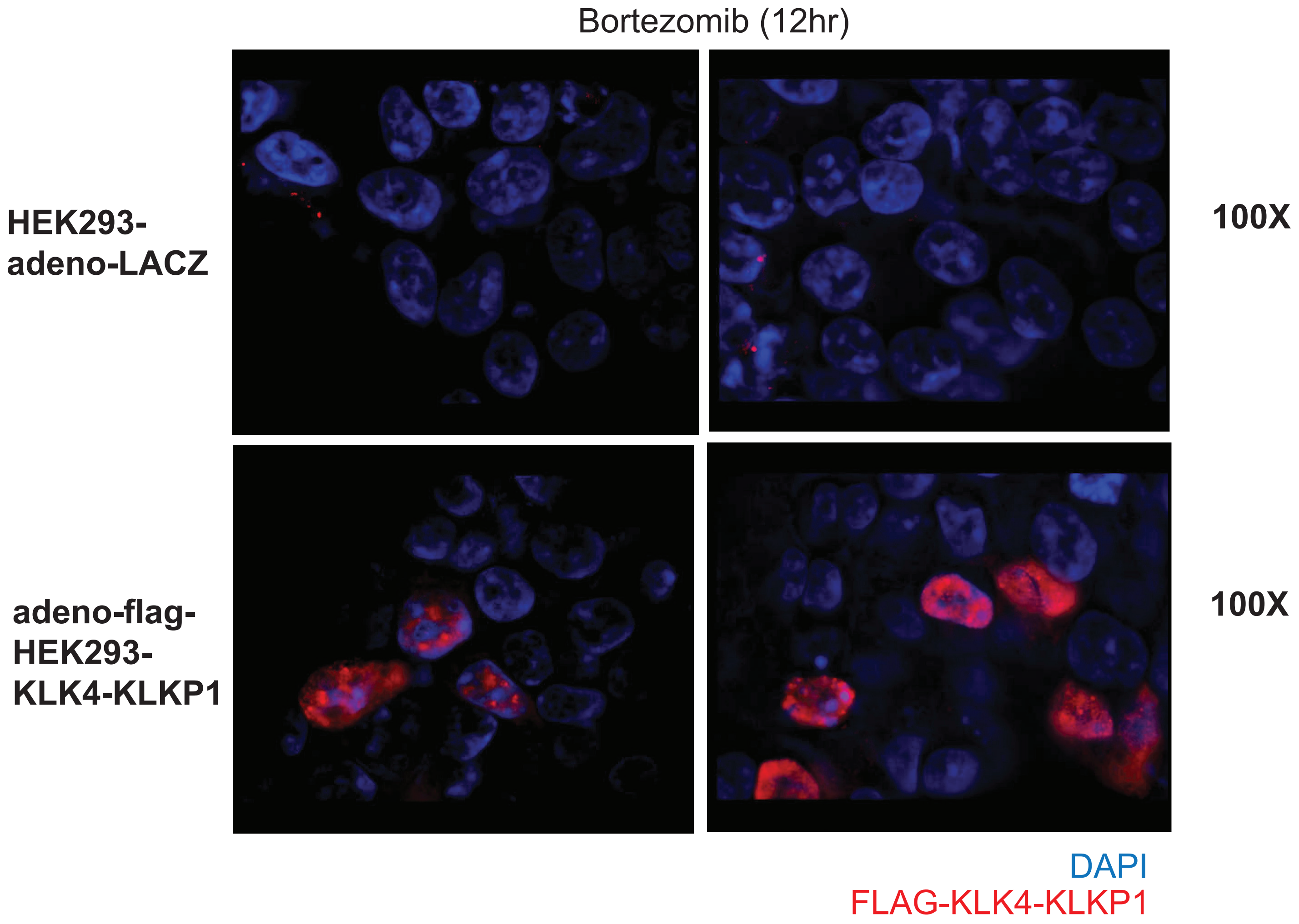


**Figure S9** : Immunofluorescence analysis of KLK4-KLKP1 in HEK293 cells. HEK293 cells were transfected with adenoviral construct carrying N-flag tagged KLK4-KLKP1 (adeno-N-Flag-KLK4-KLKP1 and treated with bortezomib for 12 hours. As a control cells transfected with adenoviral construct with LacZ (adeno-LACZ) was used. Imunofluorescence was carried out with anti-flag antibody (red) and DAPI (blue).

### Figure S10

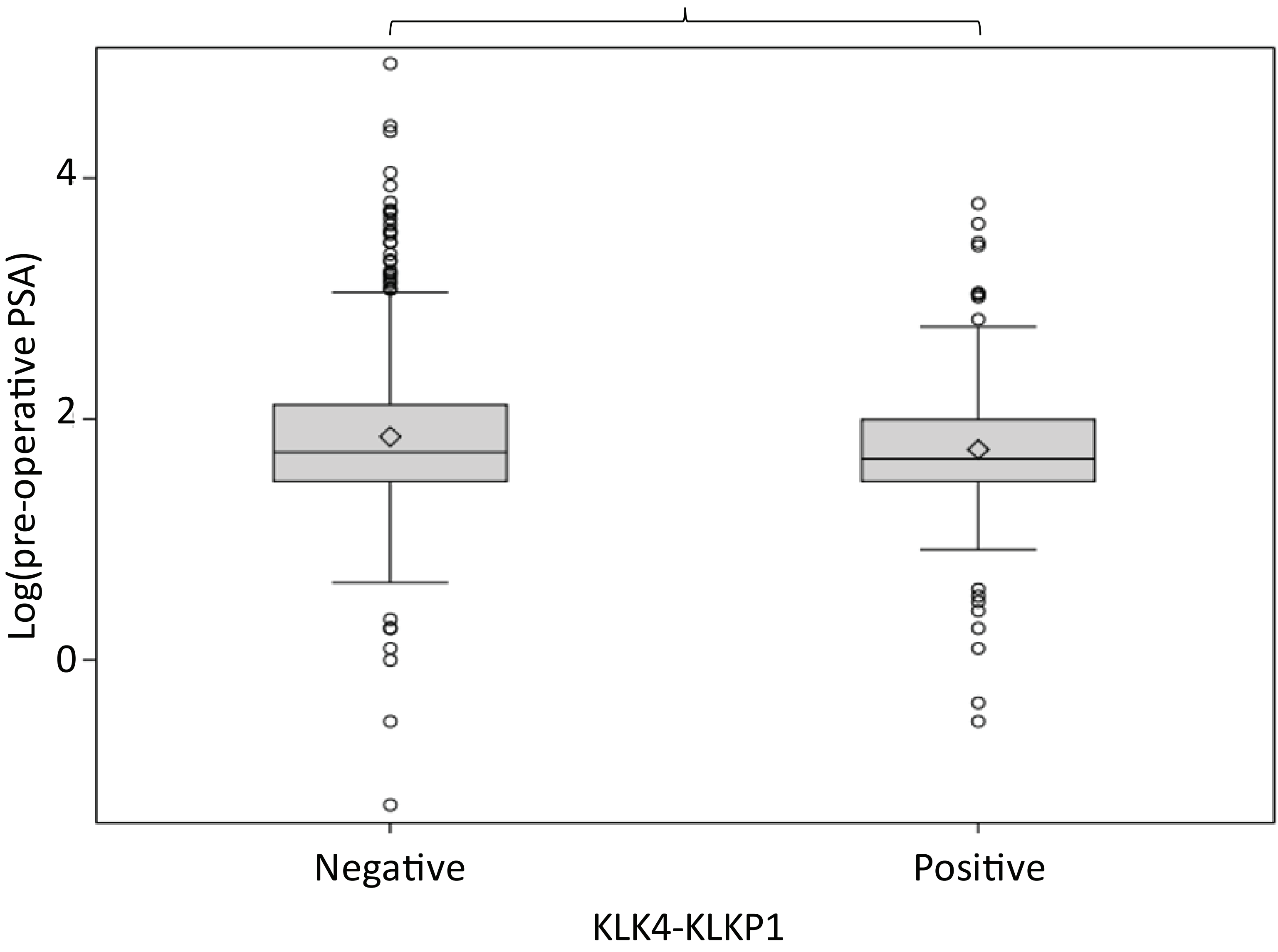


**Figure S10**: The difference in log pre-operative PSA values between KLK4-KLKP1 negative and positive cases. P value was calculated from t-test.

### Figure S11

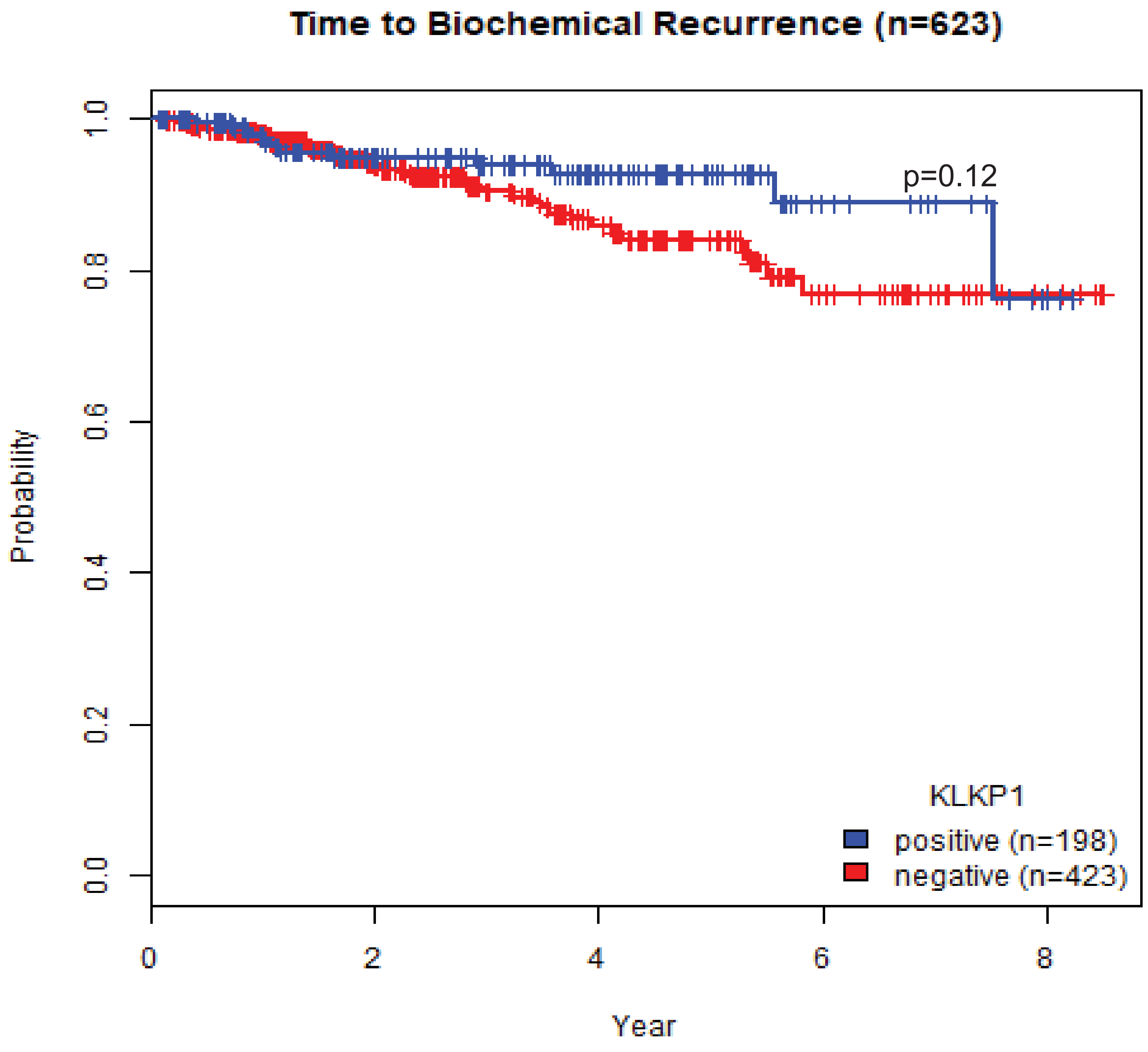


**Figure S11:** The difference in time to biochemical recurrence between KLK4-KLKP1 positive and negative patients.

### Figure S12

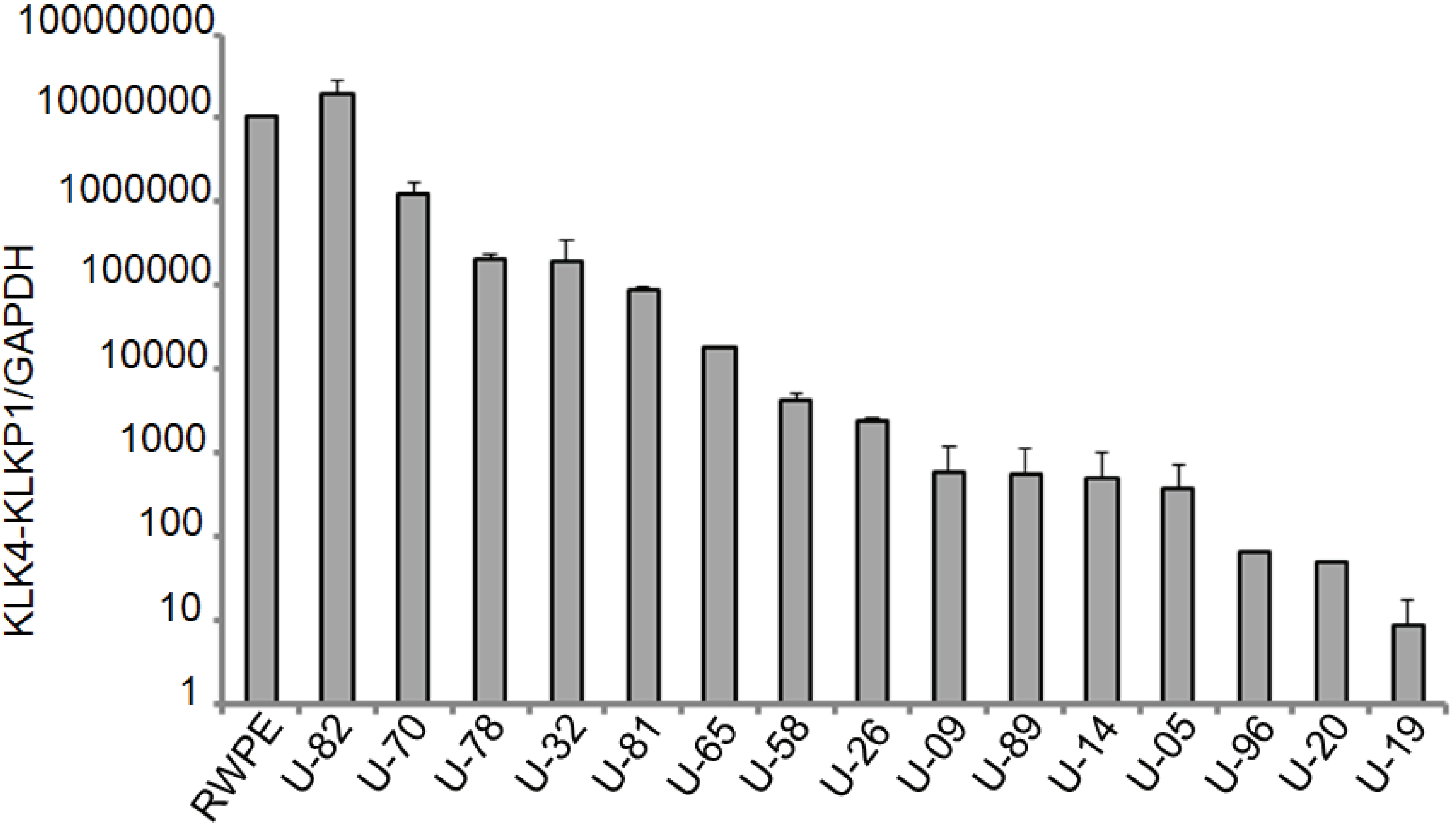


**Figure S12:** The Q-PCR analysis of KLK4-KLKP1 in urine samples of patients. Random urine samples were obtained from patients. The RNA was isolated and Q-PCR was carried out after cDNA synthesis. Q-PCR analysis was also carried out in RWPE-1 cells as a control. The number that starts with U in the figure refers to the sample number. Only the cases with detectable KLK4-KLKP1 levels are shown.
