## Supplementary material for "Pseudogene associated recurrent gene fusion in prostate cancer": Suppl methods

### SUPPLEMENTARY MATERIALS AND METHODS

#### Immunohistochemistry (IHC) and RNA-ISH for ERG, SPINK1, ETV1, ETV4 and ETV5

Staining for ERG, SPINK1, ETV1 and ETV4 was carried out using a dual RNA-ISH and a dual IHC procedure. Here, RNA-ISH was carried out first using RNAscope® 2.5 HD Duplex Reagent Kit (ACDBio, catalog #322430) per manufacturer's instructions using RNA probes specific to ETV1 (ACDBio, catalog #311411) and ETV4 (ACDBio, catalog #478571-C2). All steps were performed according to the manufacturer's instructions, except for using Betazoid DAB (1 drop DAB to 1ml Buffer, Biocare Medical, BDB2004L) instead of the green color reagent provided with the kit. Then slides were washed twice in distilled water, and once with 1X EnVision FLEX Wash Buffer (Agilent K800621-2) for 5 minutes. Peroxidazed 1 (Biocare Medical, PX968M) and Background Punisher (Biocare Medical, BP974L) was added for 5 minutes and 10 minutes respectively. Between each step, the slides were washed with 1X EnVision FLEX Wash Buffer for 5 minutes. The slides were then incubated with anti-ERG rabbit primary antibody (1:50; Abcam, ab92513) and anti-SPINK1 mouse antibody (1:100; Novus Biologicals, H00006690-M01) overnight at 4° C in a humidifying chamber, coverslipped in parafilm. The next day, slides were washed in 1X EnVision Wash Buffer for 5 minutes. Then the slides were let to sit in Mach2 Doublestain 1 (Biocare Medical, MRCT523L) for 30 minutes at room temperature in a humidifying chamber. After rinsing the slides in 1X EnVision Wash Buffer 3 times for 5 minutes each, Ferangi Blue solution (1 drop to 2.5ml buffer; Biocare Medical, FB813S) was added and slides were incubated for 7 minutes. Then slides were washed in 1X EnVision FLEX Wash Buffer for 5 minutes, and were treated with Vina Green solution (1 drop to 1ml buffer; Biocare Medical, BRR807AS) for 15 minutes. Next slides were washed 2 times in distilled water, and were incubated with EnVision FLEX Hematoxylin (Agilent, K800821-2) for 2 minutes. After rinsing the slides several times in tap water, and drying, slides were dipped in xylene approximately 15 times. Then the slides were mounted in EcoMount and scanned using the Aperio image scanner. For ETV5, staining was carried out using RNA-ISH procedure as same as for KLK4-KLKP1 RNA-ISH with RNA probes specific for ETV5 (ACDBio, 590371). ERG, SPINK1, ETV1, ETV4, and ETV5 staining were reviewed and scored as positive or negative. With ERG IHC, diffused nuclear staining was considered as positive signal for ERG over-expression. With IHC carried out for SPINK1, diffused cytoplasmic staining was defined as positive for SPINK1 over-expression. With ETV1, ETV4 and ETV5, a staining pattern of distinct punctuate cytoplasmic dots was regarded as positive for over-expression. In all cases, no visible staining pattern was considered as negative.

**RNA Isolation and cDNA Synthesis**

Total RNA was isolated from harvested cultured cells and frozen xenograft tissues using Qiazol and miRNeasy Kit (Qiagen, Maryland, USA) according to the manufacturer’s instructions. RNA integrity was verified on an Agilent Bioanalyzer 2100 (Agilent Technologies, Palo Alto, CA).

cDNA was synthesized from total RNA using Superscript III (Invitrogen) and random primers (Invitrogen).

**Quantitative Real-Time PCR**

Quantitative real-time PCR (qPCR) was performed using Taqman or SYBR green-based assays (Applied Biosystems, Foster City, CA) on a 7900HT Real-Time PCR System (Applied Biosystems) or a QuantStudio 6 Flex Real-Time PCR System (Thermo Fisher Scientific) according to standard protocols. KLK4-KLKP1 fusion specific primers were used (Forward primer sequence= 5'-ACGACCTCATGCTCATCAAGTT-3', reverse primer sequence = 5'-AGTGAGCACCCAGTGAGGAT-3'). The housekeeping gene *GAPDH* was used as a loading control. Fold changes were calculated relative to *GAPDH* and normalized to the median value of the benign samples.

**Immuohistochemistry staining of xenograft tissues for KLK4-KLKP1**

Slides with PDX tissues were provided Dr. Nora Navonne at MD Anderson Cancer Center. First slides were baked at 60° C for 2 hours.  Then slides were incubated in EnVision FLEX Target Retrieval Solution, high pH (Agilent DAKO, K800421-2) in a PT Link instrument (Agilent DAKO, PT200) at 75ºC where the slides were heated to 97ºC for 20 minutes, and then cooled to 75ºC.  After being washed in 1X EnVision FLEX Wash Buffer (Agilent DAKO, K800721-2) for 5 minutes,  the slides were treated with Peroxidazed 1 (Biocare Medical, PX968M) for 5 minutes and Background Punisher (Biocare Medical, BP974L) for 10 minutes with a wash of 1X EnVision FLEX Wash Buffer for 5 minutes between each step.  Then rabbit polyclonal KRSP1 (Eurogentec) diluted 1:50 in EnVision FLEX Antibody Diluent (Agilent DAKO, K800621-2) was added to each slide. The slides were then cover slipped with parafilm and were incubated overnight in a humidifying chamber at 4º C. Then the slides were washed in 1X EnVision Wash Buffer for 5 minutes and then incubated in Mach2 Doublestain 1 (Biocare Medical, MRCT523L) for 15 minutes at room temperature in a humidifying chamber. After rinsing the slides in 1X EnVision Wash Buffer 3 times for 5 minutes each, a Betazoid DAB solution (1 drop to 1ml buffer; Biocare Medical, BDB2004L) was added and the slides were incubated for 5 minutes. Rest of the steps were performed similar to ERG and SPINK1 IHC described above. After scanning the slides using Aperio scanner, slides were reviewed for a diffused staining pattern indicating KLK4-KLKP1 expression.

**Immunofluorescence microscopy**

The RWPE-1 cells transfected with KLK4-KLKP1 construct were plated on 18-mm cover glasses at a density of $2x10^5$ cells/well. Next day the cells were treated with Bortezomib (1µM) and incubated for 12 h. The cells were then washed with DPBS and fixed in an acetic acid: ethanol (2:1) solution for 5 min. Nonspecific binding was blocked with 5% goat and horse serum/PBS for 1 h at room temperature, and the cells were then incubated overnight with the N-flag-tag primary antibody (1:50) in a humidified chamber. After washing twice in PBS, the cells were incubated with fluorescence-labeled secondary antibody (1:100) for 1 h at room temperature in the dark. The nuclei were stained with DAPI in the dark for 30 min at room temperature. The slides were washed twice with PBS, covered with DABCO (Sigma-Aldrich) and examined with confocal laser scanning microscopy.

**Patient Derive Xenografts (PDX)**

MDA PCa PDXs were developed in the laboratory of Dr Navone at the “Prostate Cancer Patient Derived Xenografts Program”-the department of Genitourinary Medical Oncology-MD Anderson

Cancer Center and the David H. Koch Center for Applied Research of Genitourinary Cancers. PDXs were established following previously described procedures (1, 2) and propagated as subcutaneous xenografts in 6- to 8-week-old male CB17 SCID mice (Charles River Laboratories, Wilmington, MA). PDX development and expansion were conducted in accordance with accepted standards of animal care and were approved by the Institutional Animal Care and Use Committee of MD Anderson. PDXs were developed with the support of the Prostate Cancer Foundation, the David H. Koch Center for Applied Research in Genitourinary Cancers at MD Anderson and Cancer Center Prostate Cancer SPORE (NIH/NCI P50 CA140388-08).

1. Navone N, and Labanca E. In: Wang Y, Lin D, and Gout PW eds. Patient-Derived Xenograft Models of Human Cancer. Humana Press; 2017:93-114.

2. Navone NM, van Weerden WM, Vessella RL, Williams ED, Wang Y, Isaacs JT, et al. Movember GAP1 PDX project: An international collection of serially transplantable prostate cancer patient-derived xenograft (PDX) models. Prostate. 2018;78(16):1262-82.
