## Supplementary material for "Pseudogene associated recurrent gene fusion in prostate cancer": Tables 1-6

| Tumor grade | No of cases with KLK4-KLKP1 RNA-ISH negative | No of cases with KLK4-KLKP1 RNA-ISH positive | p-value |
|---|---|---|---|
| Non-cancer (Benign,HGPIN, Atypical,stroma) | 39 (83%) | 8 (17%) | |
| GG 1,2,3,4,5 | 409 (67%) | 201 (33%) | 0.02 |

**Table 1:** The comparison of KLK4-KLKP1 RNA-ISH status between non-cancer (benign, HGPIN, atypical, stroma) and GG 1-5 cases. GG refers to Gleason grade. The number of cases in each tumor grade with KLK4-KLKP1 RNA-ISH signal positive (1+ to 4+) and KLK4-KLKP1 negative is shown. The p value was calculated using Pearson's chi-square test. $p<0.05$ was considered statistically significant.

| Tumor grade | No of cases with KLK4-KLKP1 RNA-ISH negative | No of cases with KLK4-KLKP1 RNA-ISH positive | p-value |
|---|---|---|---|
| GG 1 | 70 | 39 | >0.05 |
| GG 2 | 165 | 82 | >0.05 |
| GG 3 | 79 | 39 | >0.05 |
| GG 4 | 64 | 30 | >0.05 |
| GG 5 | 31 | 11 | >0.05 |

**Table 2:** KLK4-KLKP1 RNA-ISH status compared between different Gleason grade groups. GG refers to Gleason grade. The number of KLK4-KLKP1 RNA-ISH signal positive (1+ to 4+) cases in each Gleason grade group is shown. The association of KLK1-KLKP1 RNA-ISH signal with Gleason grade was analyzed using Pearson's chi-square test. $p<0.05$ was considered statistically significant.

| Race | No of cases with KLK4-KLKP1 RNA-ISH positive | No of cases with KLK4-KLKP1 RNA-ISH negative | p-value |
|---|---|---|---|
| African American (AA) | 69 (28%) | 181 (72%) | |
| Caucasian American (CA) | 128 (34%) | 250(66%) | 0.098 |

**Table 3:** KLK4-KLKP1 RNA-ISH status compared between AA and CA patients. The number of cases with KLK4-KLKP1 RNA-ISH signal positive (1+ to 4+) and KLK4-KLKP1 negative in each race group are shown. The p value was calculated using Pearson's chi-square test. $p<0.05$ was considered statistically significant.

| Age group | No of cases with KLK4-KLKP1 RNA-ISH positive | No of cases with KLK4-KLKP1 RNA-ISH negative | p-value |
|---|---|---|---|
| Young age (less than 50 years) | 26 (57%) | 20 (43%) | |
| Old age (equal to 50 years or higher) | 183 (30%) | 427 (70%) | 0.0002 |

**Table 4:** KLK4-KLKP1 RNA-ISH status compared between young (age lower than 50 years) and old (age equal to or higher than 50 years) patients. The number of cases with KLK4-KLKP1 RNA-ISH signal positive (1+ to 4+) and KLK4-KLKP1 negative in each age group are shown. The p value was calculated using Pearson's chi-square test. $p<0.05$ was considered statistically significant.

| Molecular Marker | No of cases with the corresponding marker status | No of cases with KLK4-KLKP1 RNA-ISH positive | No of cases with KLK4-KLKP1 RNA-ISH negative | p value |
|---|---|---|---|---|
| ERG | No of cases with ERG negative | 130 (63%) | 355 (80%) | |
| | No of cases with ERG positive | 78 (38%) | 90 (20%) | <.001 |
| SPINK1 | No of cases with SPINK1 negative | 184 (88%) | 387 (87%) | |
| | No of cases with SPINK1 positive | 24 (12%) | 57 (13%) | 0.703 |
| ETV1 | No of cases with ETV1 negative | 197 (95%) | 423 (95%) | |
| | No of cases with ETV1 positive | 11 (5%) | 22 (5%) | 0.849 |
| ETV4 | No of cases with ETV4 negative | 200 (96%) | 439 (99%) | |
| | No of cases with ETV4 positive | 8 (4%) | 6 (1%) | 0.077 |
| ETV5 | No of cases with ETV5 negative | 193 (92%) | 425 (95%) | 0.217 |
| | No of cases with ETV5 positive | 16 (8%) | 23 (5%) | |
| PTEN | No of cases with PTEN loss | 57 (28%) | 159 (36%) | 0.032 |
| | No of cases without PTEN loss | 150 (72%) | 281 (64%) | |

**Table 5:** ERG, SPINK1, ETV1, ETV4 and ETV5 marker status compared with KLK4-KLKP1 status. The number of cases with KLK4-KLKP1 RNA-ISH signal positive (1+ to 4+) and KLK4-KLKP1 negative in each marker status is shown. The p value was calculated using Pearson's chi-square test. $p<0.05$ was considered statistically significant.

**Table 6**

| MDA PCa PDXs and a cell line | Source | Treatment |
|---|---|---|
| MDA PCa 173-13 | Testis | Therapy naïve |
| MDA PCa 149-1 | Bladder, local extension of prostate cancer | CRPC |
| MDA PCa 153-7 | Thyroid | CRPC |
| MDA PCa 2b | Bone | CRPC |
| MDA PCa 183-A | Bone | Therapy naïve |
| MDA PCa 203-A | Bone | CRPC |
| MDA PCa 101 | Liver | CRPC |
| MDA PCa 152-1 | Brain | CRPC |
| MDA PCa 175-10 | Testis | CRPC |
| MDA PCa 180-30 | Prostate | CRPC |
| MDA PCa 153-14 | Thyroid | CRPC |
| MDA PCa 175-2 | Testis | CRPC |
| MDA PCa 230-A | Chest wall | CRPC |
| MDA PCa 146-12 | Bladder, local extension of prostate cancer | CRPC |
| MDA PCa 188-2 | Bladder, local extension of prostate cancer | CRPC |
| MDA PCa 133-4 | Bone | CRPC |
| MDA PCa 180-11 | Bladder, local extension of prostate cancer | CRPC |
| MDA PCa 273-A | Retropeitoneal LN | CRPC |
| MDA PCa 94 | Pleural effusion | CRPC |
| MDA PCa 118a | Bone | CRPC |
| MDA PCa 144-4 | Prostate | CRPC |
| MDA PCa 144-13 | Bladder, local extension of prostate cancer | CRPC |
| MDA PCa 146-10 | Bladder, local extension of prostate cancer | CRPC |
| MDA PCa 146-20 | Bladder, local extension of prostate cancer | CRPC |
| MDA PCa 150-1 | Bone | CRPC |
| MDA PCa 155-12 | Bladder, local extension of prostate cancer | CRPC |
| MDA PCa 155-16 | Prostate | CRPC |
| MDA PCa 166-8 | Bladder, local extension of prostate cancer | CRPC |
| MDA PCa 177-B | Prostate | CRPC |
| MDA PCa 118b | Bone | CRPC |

**Table 6:** List of PDX models used to screen KLK4-KLKP1 fusion gene.
